# Development of a Synthetic Poxvirus-Based SARS-CoV-2 Vaccine

**DOI:** 10.1101/2020.07.01.183236

**Authors:** Flavia Chiuppesi, Marcela d’Alincourt Salazar, Heidi Contreras, Vu H Nguyen, Joy Martinez, Soojin Park, Jenny Nguyen, Mindy Kha, Angelina Iniguez, Qiao Zhou, Teodora Kaltcheva, Roman Levytskyy, Nancy D Ebelt, Tae Hyuk Kang, Xiwei Wu, Thomas Rogers, Edwin R Manuel, Yuriy Shostak, Don J Diamond, Felix Wussow

**Author notes:** co-corresponding and co-senior authors. **One sentence summary:** Chiuppesi *et al.* demonstrate the use of a uniquely designed and fully synthetic poxvirus-based vaccine platform to rapidly develop a SARS-CoV-2 vaccine candidate enabling stimulation of potent humoral and cellular immune responses to multiple antigens.

## Abstract

Modified Vaccinia Ankara (MVA) is a highly attenuated poxvirus vector that is widely used to develop vaccines for infectious diseases and cancer. We developed a novel vaccine platform based on a unique three-plasmid system to efficiently generate recombinant MVA vectors from chemically synthesized DNA. In response to the ongoing global pandemic caused by SARS coronavirus-2 (SARS-CoV-2), we used this novel vaccine platform to rapidly produce fully synthetic MVA (sMVA) vectors co-expressing SARS-CoV-2 spike and nucleocapsid antigens, two immunodominant antigens implicated in protective immunity. Mice immunized with these sMVA vectors developed robust SARS-CoV-2 antigen-specific humoral and cellular immune responses, including potent neutralizing antibodies. These results demonstrate the potential of a novel vaccine platform based on synthetic DNA to efficiently generate recombinant MVA vectors and to rapidly develop a multi-antigenic poxvirus-based SARS-CoV-2 vaccine candidate.

## Introduction

Modified Vaccinia Ankara (MVA) is a highly attenuated poxvirus vector that is widely used to develop vaccine approaches for infectious diseases and cancer (*1–3*). As a result of the attenuation process through 570 virus passages on chicken embryo fibroblast (CEF), MVA has acquired multiple major and minor genome alterations (*4, 5*), leading to severely restricted host cell tropism (*6*). MVA can efficiently propagate on CEF and a baby hamster kidney (BHK) cell line, while in most mammalian cells, including human cells, MVA replication is limited due to a late block in virus assembly (*3, 6*). Its excellent safety and immunogenicity profile in addition to its versatile expression system and large capacity to incorporate heterologous DNA make MVA an ideal vector for recombinant vaccine development (*1, 7*). We developed MVA vaccines for animal models of cytomegalovirus-associated disease in pregnant women while demonstrating vaccine efficacy in several clinical trials in solid tumor and stem cell transplant patients (*8–13*).

Since the outbreak of the novel severe acute respiratory syndrome coronavirus-2 (SARS-CoV-2) in December 2019 (*14, 15*), the virus has spread to more than 200 countries worldwide, causing a pandemic of global magnitude with over 400,000 deaths. Many vaccine candidates are currently under rapid development to control this global pandemic (*16–18*), some of which have entered into clinical trials with unprecedented pace (*17, 19*). Most of these approaches employ antigenic forms of the Spike (S) protein as it is considered the primary target of protective immunity (*16, 20–22*). The S protein mediates SARS-CoV-2 entry into a host cell through binding to angiotensin-converting enzyme 2 (ACE) and is the major target of neutralizing antibodies (NAb) (*23–25*). Studies in rhesus macaques show that vaccine strategies based on the S antigen can prevent SARS-CoV-2 infection and disease in this relevant animal model (*18*), indicating that the S antigen may be sufficient as a vaccine immunogen to elicit SARS-CoV-2 protective immunity. However, a recent study demonstrated that even patients without measurable NAb can recover from SARS-CoV-2 infection, suggesting that protection against SARS-CoV-2 infection is mediated by both humoral and cellular immunity to multiple immunodominant antigens, including S and nucleocapsid (N) antigens (*20, 26*).

We developed a novel vaccine platform based on a uniquely designed three-plasmid system to efficiently generate recombinant MVA vectors from chemically synthesized DNA. In response to the ongoing global pandemic caused by SARS-CoV-2, we used this novel vaccine platform to rapidly produce synthetic MVA (sMVA) vectors co-expressing full-length S and N antigens. We demonstrate that these sMVA vectors stimulate robust SARS-CoV-2 antigen-specific humoral and cellular immunity in mice, including potent NAb. These results emphasize the value of a novel vaccine platform based on synthetic DNA to efficiently produce recombinant poxvirus vectors and warrant further pre-clinical and clinical testing of a multi-antigenic sMVA vaccine candidate to control the ongoing SARS-CoV-2 pandemic and its devastating consequences.

## Results

### Construction of sMVA

To develop the three-plasmid system of the sMVA vaccine platform, we designed three unique synthetic sub-genomic MVA fragments (sMVA F1-F3) based on the MVA genome sequence published by Antoine *et al.* (*4*), which is ~178 kbp in length and contains ~9.6 kbp inverted terminal repeats (ITRs) (Figure 1A). The three fragments were designed as follows: sMVA F1 comprises ~60 kbp of the left part of the MVA genome, including the left ITR sequences; sMVA F2 contains ~60 kbp of the central part of the MVA genome; and sMVA F3 contains ~60 kbp of the right part of the MVA genome, including the right ITR sequences (Figure 1B). sMVA F1 and F2 as well as sMVA F2 and F3 were designed to share ~3kb overlapping homologous sequences to promote recombination of the three sMVA fragments (Figure 1B). In addition, a duplex copy of the 165-nucleotide long MVA terminal hairpin loop (HL) flanked by concatemeric resolution (CR) sequences was added to both ends of each of the three sMVA fragments (Figure 1C). Such CR/HL/CR sequence arrangements are formed at the genomic junctions in poxvirus DNA replication intermediates and are essential for genome resolution and packaging (*27–31*). When circular DNA plasmids containing these CR/HL/CR sequence arrangements are transfected into helper virus-infected cells they spontaneously resolve into linear minichromosomes with intact terminal HL sequences (*28, 29, 32*). Based on these findings, we hypothesized that the three sMVA fragments designed as shown in Figure 1B-C, when co-transfected as circular DNA plasmids into helper virus-infected cells, resolve into linear minichromosomes, recombine with each other via the shared homologous sequences, and are ultimately packaged as full-length genomes into sMVA virus particles. All three sMVA fragments were cloned in *E. coli* as bacterial artificial chromosome (BAC) clones.

**Figure 1.**
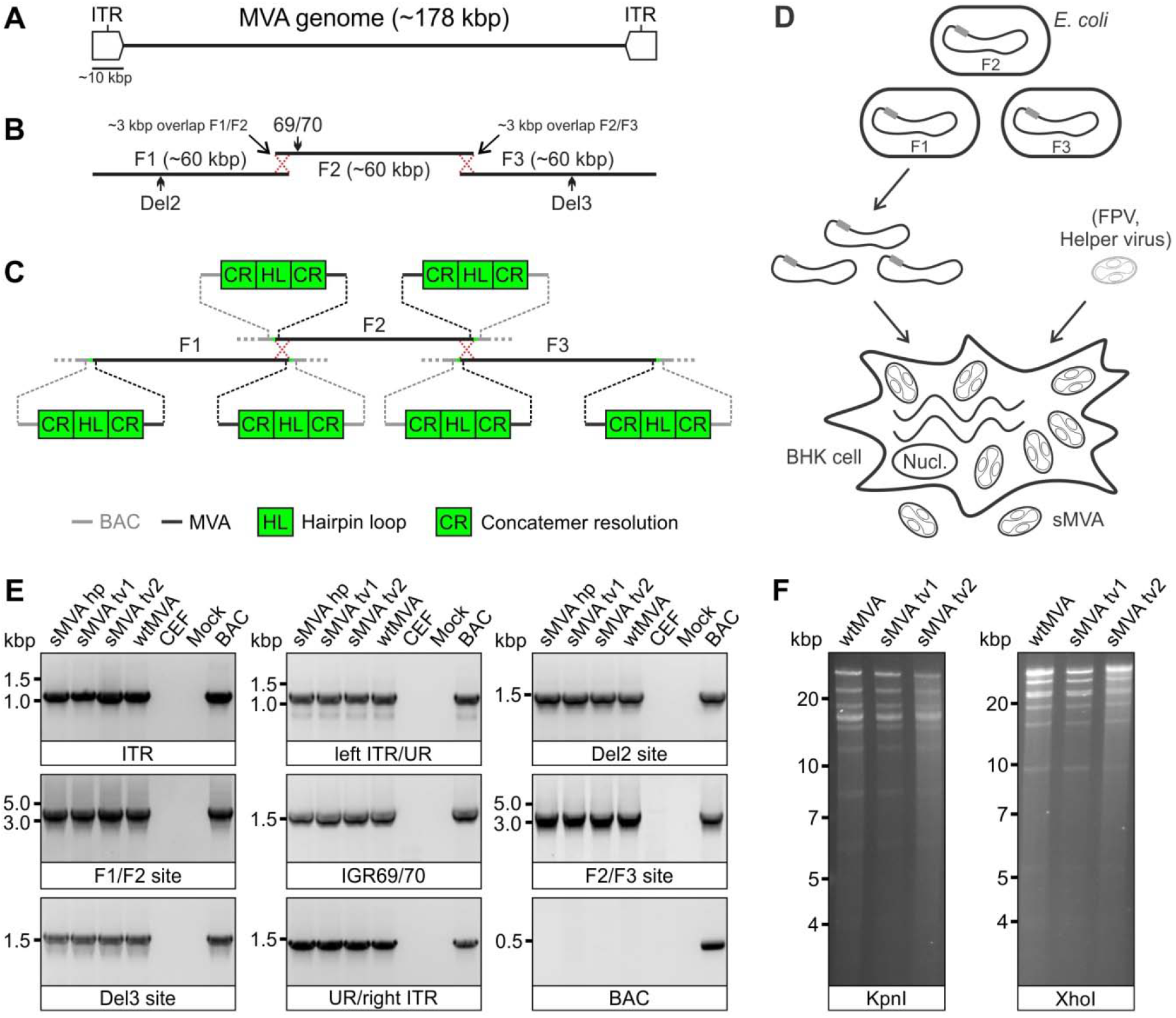
sMVA construction and characterization. **A)** Schematic of MVA genome. The MVA genome is ~178 kbp in length and contains ~9.6 kbp inverted terminal repeat (ITR) sequences. **B)** sMVA fragments. The three sub-genomic sMVA fragments (F1-F3) comprise ~60 kbp of the left, central, and right part of the MVA genome as indicated. sMVA F1/F2 and F2/F3 share ~3 kbp overlapping homologous sequences for recombination (red dotted crossed lines). Approximate genome positions of commonly used MVA insertion (Del2, IGR69/70, Del3) are indicated **C)** Terminal CR/HL/CR sequences. Each of the sMVA fragments contains at both ends a sequence composition comprising a duplex copy of the MVA terminal hairpin loop (HL) flanked by concatemeric resolution (CR) sequences. BAC = bacterial artificial chromosome vector. **D)** sMVA reconstitution. The sMVA fragments are isolated from the *E. coli* and co-transfected into BHK cells, which are subsequently infected with FPV as a helper virus to initiate sMVA virus reconstitution. **E)** PCR analysis. CEF infected with sMVA, derived with FPV HP1.441 (sMVA hp) or TROVAC from two independent virus reconstitutions (sMVA tv1 and sMVA tv2), were investigated by PCR for several MVA genome positions (ITR sequences, transition left or right ITR into internal unique region (left ITR/UR; UR/right ITR), Del2, IGR69/70 and Del3 insertion sites, and F1/F2 and F2/F3 recombination sites) and absence of BAC vector sequences. PCR reactions with wtMVA-infected and uninfected cells, without sample (mock), or with MVA BAC were performed as controls. **F)** Restriction fragment length analysis. Viral DNA isolated from ultra-purified sMVA (sMVA tv1 and sMVA tv2) or wtMVA virus was compared by KpnI and XhoI restriction enzyme digestion.

Using a previously employed procedure to rescue MVA from a BAC (*8, 9, 33*), sMVA virus was reconstituted with Fowl pox (FPV) as a helper virus upon co-transfection of the three DNA plasmids into BHK cells (Figure 1D), which are non-permissive for FPV(*34*). Two different FPV strains (HP1.441 and TROVAC) (*35, 36*) were used to promote sMVA virus reconstitution (Figure 2A). Ultra-purified sMVA virus was produced following virus propagation in CEF, which are commonly used for MVA vaccine production. The virus titers achieved with reconstituted sMVA virus were similar to virus titers achieved with “wild-type” MVA (wtMVA) (Table S1).

**Figure 2.**
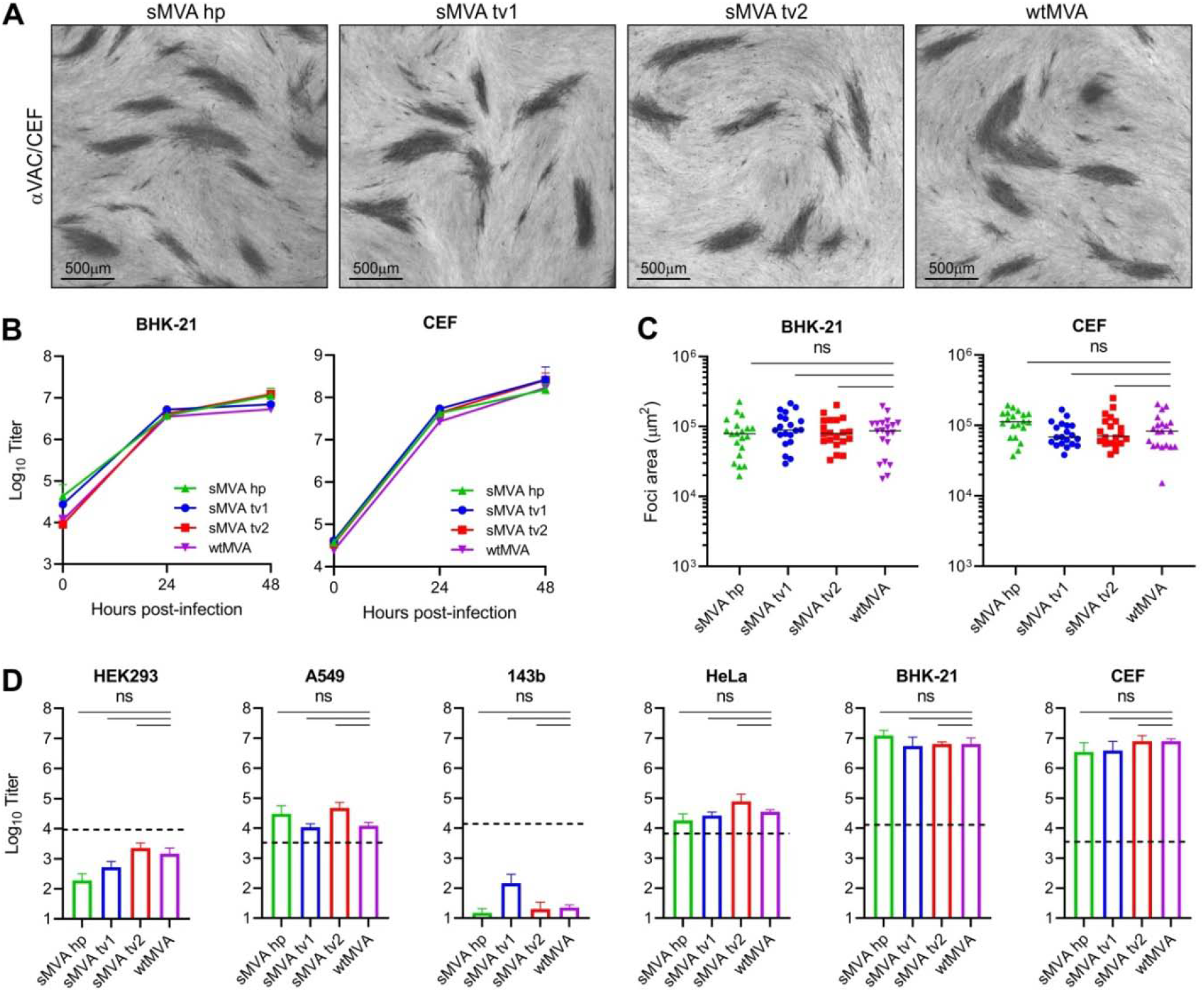
sMVA replication properties. The replication properties of sMVA derived with FPV HP1.441 (sMVA hp) or TROVAC from two independent sMVA virus reconstitution (sMVA tv1 and sMVA tv2) were compared with wtMVA. **A)** Viral foci. CEF infected at low multiplicity of infection (MOI) with the reconstituted sMVA virus or wtMVA were immunostained using anti-Vaccinia polyclonal antibody (αVAC). **B)** Replication kinetics. BHK or CEF cells were infected at 0.02 MOI with sMVA or wtMVA and viral titers of the inoculum and infected cells at 24 and 48 hours post infection were determined on CEF. Mixed-effects model with the Geisser-Greenhouse correction was applied; at 24 and 48 hours post-infection differences between groups were not significant. **C)** Viral foci size analysis. BHK or CEF cell monolayers were infected at 0.002 MOI with sMVA or wtMVA and areas of viral foci were determined at 24 hours post infection following immunostaining with αVAC antibody. **D)** Host cell range analysis. Various human cell lines (HEK293, A549, 143b, and HeLa), CEF or BHK cells were infected at 0.01 MOI with sMVA or wtMVA and virus titers were determined at 48 hours post infection on CEF. Dotted lines indicate the calculated virus titer of the inoculum based on 0.01 MOI. Differences between groups in C-D were calculated using one-way ANOVA followed by Tukey’s (C) or Dunnett’s (D) multiple comparison tests. ns = not significant.

### *In vitro* characterization of sMVA

To characterize the viral DNA of sMVA, DNA extracts from sMVA and wtMVA-infected CEF were compared for several MVA genome positions by PCR. Similar PCR results were obtained with sMVA and wtMVA for all evaluated genome positions (Figure 1E), including the F1/F2 and F2/F3 recombination sites, indicating efficient recombination of the three sMVA fragments. Additional PCR analysis indicated the absence of any BAC vector sequences in sMVA viral DNA (Figure 1E), suggesting spontaneous and efficient removal of bacterial vector elements upon sMVA virus reconstitution. Comparison of viral DNA from ultra-purified sMVA and wtMVA virus by restriction enzyme digestion revealed similar genome pattern between sMVA and wtMVA (Figure 1F). Sequencing analysis of the sMVA viral DNA confirmed the MVA genome sequence at several positions, including the F1/F2 and F2/F3 recombination sites. Furthermore, whole genome sequencing analysis of one of the sMVA virus isolates reconstituted with FPV TROVAC confirmed the assembly of the reference MVA genome sequence and absence of vector-specific sequences in viral DNA originating from reconstituted sMVA virus.

To characterize the replication properties of sMVA, growth kinetics of sMVA and wtMVA were compared on BHK and CEF cells, two cell types known to support productive MVA replication (*6*). This analysis revealed similar growth kinetics of sMVA and wtMVA on both BHK and CEF cells (Figure 2B). In addition, similar areas of viral foci were determined in BHK and CEF cell monolayers infected with sMVA or wtMVA (Figure 2C), suggesting similar capacity of sMVA and wtMVA to spread in MVA permissive cells. Compared to the productive replication of sMVA and wtMVA in BHK and CEF cells (*6*), only limited virus production was observed with sMVA or wtMVA following infection of various human cell lines (Figure 2D). These results are consistent with the severely restricted replication properties of MVA and show that the sMVA virus can efficiently propagate in BHK and CEF cells, while it is unable to propagate in human cells.

### *In vivo* immunogenicity of sMVA

To characterize sMVA *in vivo*, the immunogenicity of sMVA and wtMVA was compared in C57BL/6 mice following two immunizations at high or low dose. MVA-specific binding antibodies stimulated by sMVA and wtMVA after the first and second immunization were comparable (Figures 3A, S1A). While the antibody levels in the high dose vaccine groups exceeded those of the low dose vaccine groups after the first immunization, similar antibody levels in the high and low dose vaccine groups were observed after the second immunization. In addition, no significant differences were detected in the levels of MVA-specific NAb responses induced by sMVA and wtMVA after the second immunization (Figures 3B, S1B). MVA-specific T cell responses determined after the booster immunization by *ex vivo* antigen stimulation using immunodominant peptides (*37*) revealed similar MVA-specific T cell levels in mice receiving sMVA or wtMVA (Figures 3C-D and S1C-D). These results indicate that the sMVA virus has a similar capacity to wtMVA in inducing MVA-specific humoral and cellular immunity in mice.

**Figure 3.**
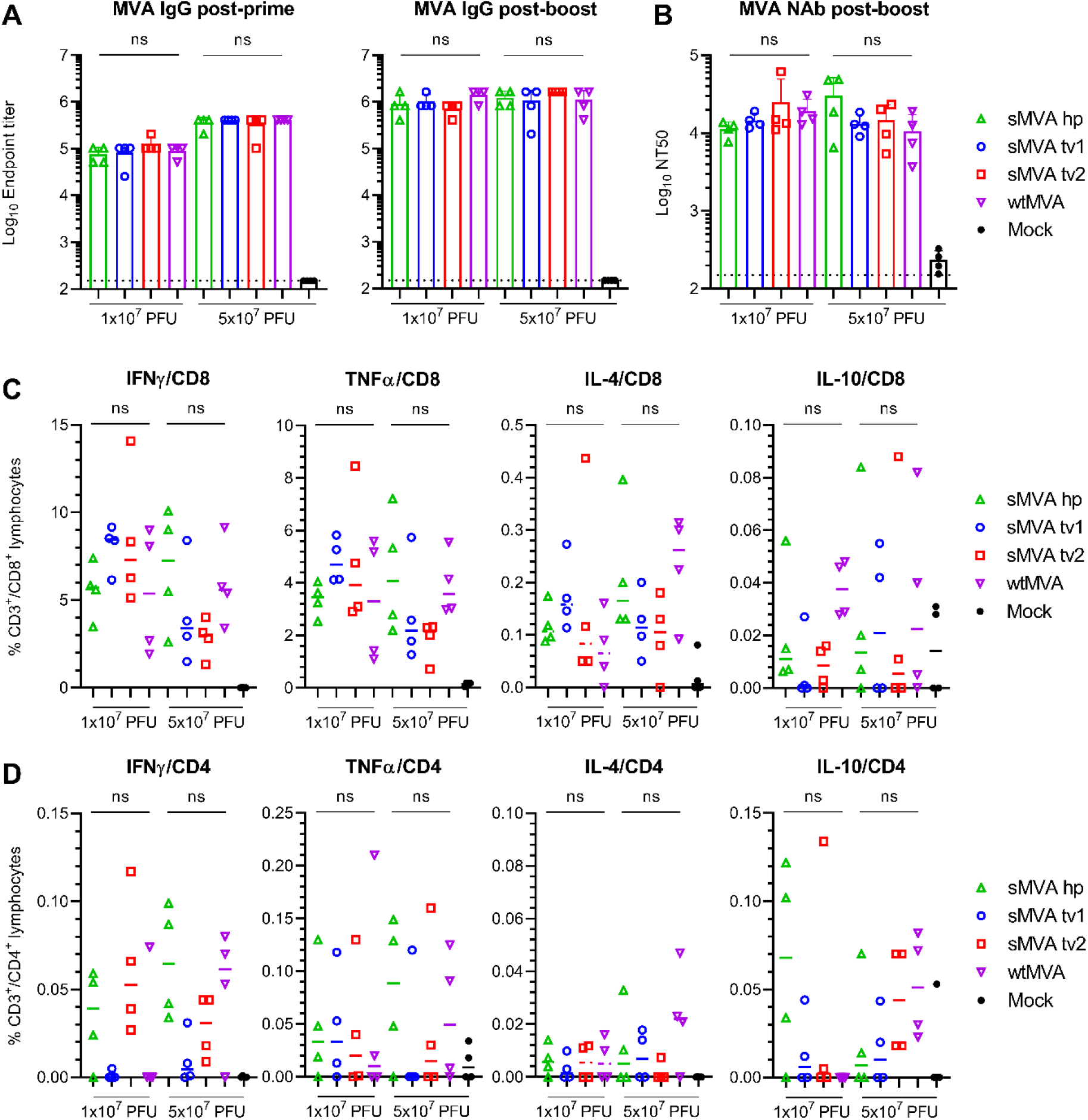
sMVA *in vivo* immunogenicity. sMVA derived either with FPV HP1.441 (sMVA hp) or TROVAC from two independent virus reconstitution (sMVA tv1 and sMVA tv2) was compared by *in vitro* analysis with wtMVA. C57BL/6 mice were immunized twice at three week interval with low (1×10^7^ PFU) or high (5×10^7^ PFU) dose of sMVA or wtMVA. Mock-immunized mice were used as controls **A)** Binding antibodies. MVA-specific binding antibodies (IgG titer) stimulated by sMVA or wtMVA were measured after the first and second immunization by ELISA. **B)** NAb responses. MVA-specific NAb titers induced by sMVA or wtMVA were measured after the booster immunization against recombinant wtMVA expressing a GFP marker. **C-D)** T cell responses. MVA-specific IFN*γ*, TNFα, IL-4, and IL-10-secreting CD8+ (C) and CD4+ (D) T cell responses induced by sMVA or wtMVA after two immunizations were measured by flow cytometry following *ex vivo* antigen stimulation using B8R immunodominant peptides. Differences between groups were evaluated using one-way ANOVA with Tukey’s multiple comparison test. ns = not significant.

### Construction of sMVA SARS-CoV-2 vaccine vectors

Using highly efficient BAC recombination techniques in *E. coli*, full-length SARS-CoV-2 S and N antigen sequences were inserted into commonly used MVA insertions sites located at different positions within the three sMVA fragments. Combinations of modified and unmodified sMVA fragments were subsequently co-transfected into FPV-infected BHK cells to reconstitute sMVA SARS-CoV-2 (sMVA-CoV2) vectors expressing the S and N antigen sequences alone or combined (Figure 4A and 4B). In the single recombinant vectors encoding S or N alone, termed sMVA-S and sMVA-N, the antigen sequences were inserted into the Deletion (Del3) site (Figures 1B and 4B) (*5*). In the double recombinant vectors encoding both S and N, termed sMVA-N/S and sMVA-S/N, the antigen sequences were inserted into Del3 and the Deletion 2 (Del2) site (sMVA-N/S), or they were inserted into Del3 and the intergenic region between 069R and 070L (IGR69/70) (sMVA-S/N) (Figures 1B and 4B) (*5, 38*). All antigen sequences were inserted into sMVA together with mH5 promoter to promote antigen expression during early and late phase of MVA replication (*39, 40*). sMVA-CoV-2 vaccine vectors were reconstituted with FPV HP1.441 or TROVAC. Ultra-purified virus of the sMVA-CoV2 vectors produced using CEF reached titers that were similar to those achieved with sMVA or wtMVA (Table S1).

**Figure 4.**
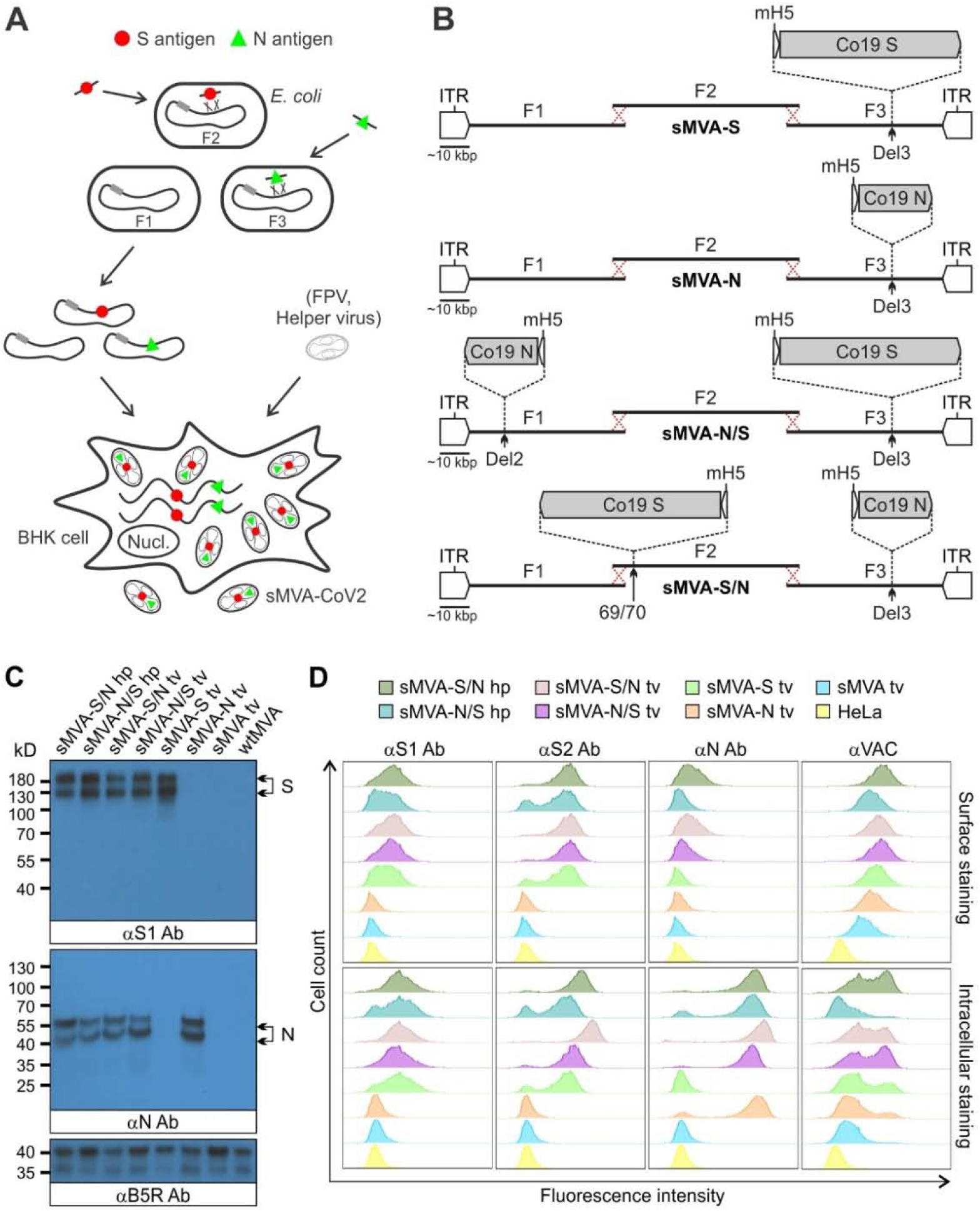
Construction and characterization of sMVA-CoV2 vectors. **A)** Schematic representation of vector construction. S and N antigen sequences (red spheres and green triangles) were inserted into sMVA fragments F2 and F3 by bacterial recombination methods in *E. coli*. The modified sMVA fragments of F1 and F2 with inserted antigen sequences and the unmodified sMVA fragment F1 were isolated from *E. coli* and co-transfected into FPV-infected BHK cells to initiate virus reconstitution. **B)** Schematics of single (sMVA-S, sMVA-N) and double (sMVA-N/S, sMVA-S/N) recombinant sMVA-CoV2 vectors with S and N antigen sequences inserted into commonly used MVA insertion sites (Del2, IGR69/70, Del3). All antigens were expressed via the Vaccinia mH5 promoter. **C)** Western Blot. BHK cells infected with the single and double recombinant sMVA-CoV2 vectors derived with FPV HP1.441 (sMVA-S/N hp, sMVA-N/S hp) or TROVAC (sMVA-S/N tv, sMVA-N/S tv, sMVA-S tv, sMVA-N tv) were evaluated for antigen expression by Western Blot using anti-S1 and N antibodies (αS1 and αN Ab). Vaccinia B5R protein was verified as infection control. Higher and lower molecular weight bands may represent mature and immature protein species. **D)** Flow cytometry staining. HeLa cells infected with the vaccine vectors were evaluated by cell surface and intracellular flow staining using anti-S1, S2, and N antibodies (αS1, αS2, and αN Ab). Live cells were used to evaluate cell surface antigen expression. Fixed and permeabilized cells were used to evaluate intracellular antigen expression. Anti-Vaccinia virus antibody (αVAC) was used as staining control to verify MVA protein expression. Cells infected with sMVA or wtMVA or uninfected cells were used as controls for experiments in C and D as indicated.

### *In vitro* characterization of sMVA-CoV2 vaccine vectors

To characterize S and N antigen expression by the sMVA-CoV2 vectors, BHK cells infected with the sMVA-CoV2 vectors were evaluated by Immunoblot using S and N-specific antibodies. This analysis confirmed the expression of the S or N antigen alone by the single recombinant vaccine vectors sMVA-S and sMVA-N, while the expression of both the S and the N antigen was confirmed for the double recombinant vectors sMVA-N/S and sMVA-S/N (Figure 4C).

Further characterization of the antigen expression by the sMVA-CoV2 vectors in HeLa cells using cell surface and intracellular flow cytometry (FC) staining confirmed single and dual S and N antigen expression by the single and double recombinant vaccine vectors. Staining with S-specific antibodies revealed abundant cell surface and intracellular antigen expression by all vectors encoding the S antigen (sMVA-S, sMVA-N/S, sMVA-S/N) (Figure 4D). In contrast, staining with anti-N antibody revealed predominantly intracellular antigen expression by all vectors encoding the N antigen (sMVA-N, sMVA-N/S, sMVA-S/N) (Figure 4D), although cell surface staining was observed to a minor extent. S and N antigen expression by the sMVA-CoV2 vectors was also investigated by immunofluorescence. This analysis confirmed co-expression of the S and N antigens by the double recombinant vaccine vectors and indicated efficient cell surface and intracellular expression of the S antigen, whereas the expression of the N antigen was predominantly observed intracellular (Figure S2A-C). These results demonstrate efficient antigen expression by the single and double recombinant sMVA-CoV2 vectors.

### *In vivo* immunogenicity of sMVA-CoV2 vectors

To determine the immunogenicity of the sMVA-vectored S and N antigens alone or combined, SARS-CoV-2-specific humoral and cellular immune responses were evaluated in Balb/c mice by two immunizations with the single or double recombinant vaccine vectors. High-titer antigen-specific binding antibodies were detected in all vaccine groups after the first immunization, and an increase in these responses was observed after the booster immunization (Figure 5A-B and S3A-B). While the single recombinant vectors induced binding antibodies only against the S or N antigen, the double recombinant vectors induced binding antibodies against both the S and N antigens. In addition, all sMVA-CoV2 vectors encoding the S antigen (sMVA-S, sMVA-S/N, sMVA-N/S) stimulated high-titer binding antibodies against the S receptor binding domain (RBD), which is considered the primary target of NAb(*22, 24*). Antigen-specific binding antibody titers between the single and double recombinant vaccine groups were comparable. Notably, SARS-CoV-2 antigen-specific binding antibody responses stimulated by the sMVA-CoV2 vaccine vectors in mice exceeded SARS-CoV-2 S-, RBD-, and N-specific binding antibody responses measured in human convalescent immune sera (Figures 5A-B, and Figure S4). Similar binding antibody responses to those induced by sMVA-CoV2 vectors in Balb/c mice were elicited by the vaccine vectors in C57BL/6 mice (Figure S5). Analysis of the IgG2a/IgG1 isotype ratio of the binding antibodies revealed Th-1-biased immune responses skewed toward IgG2a independently of the investigated vaccine group or antigen (Figure 5C and S3C).

**Figure 5.**
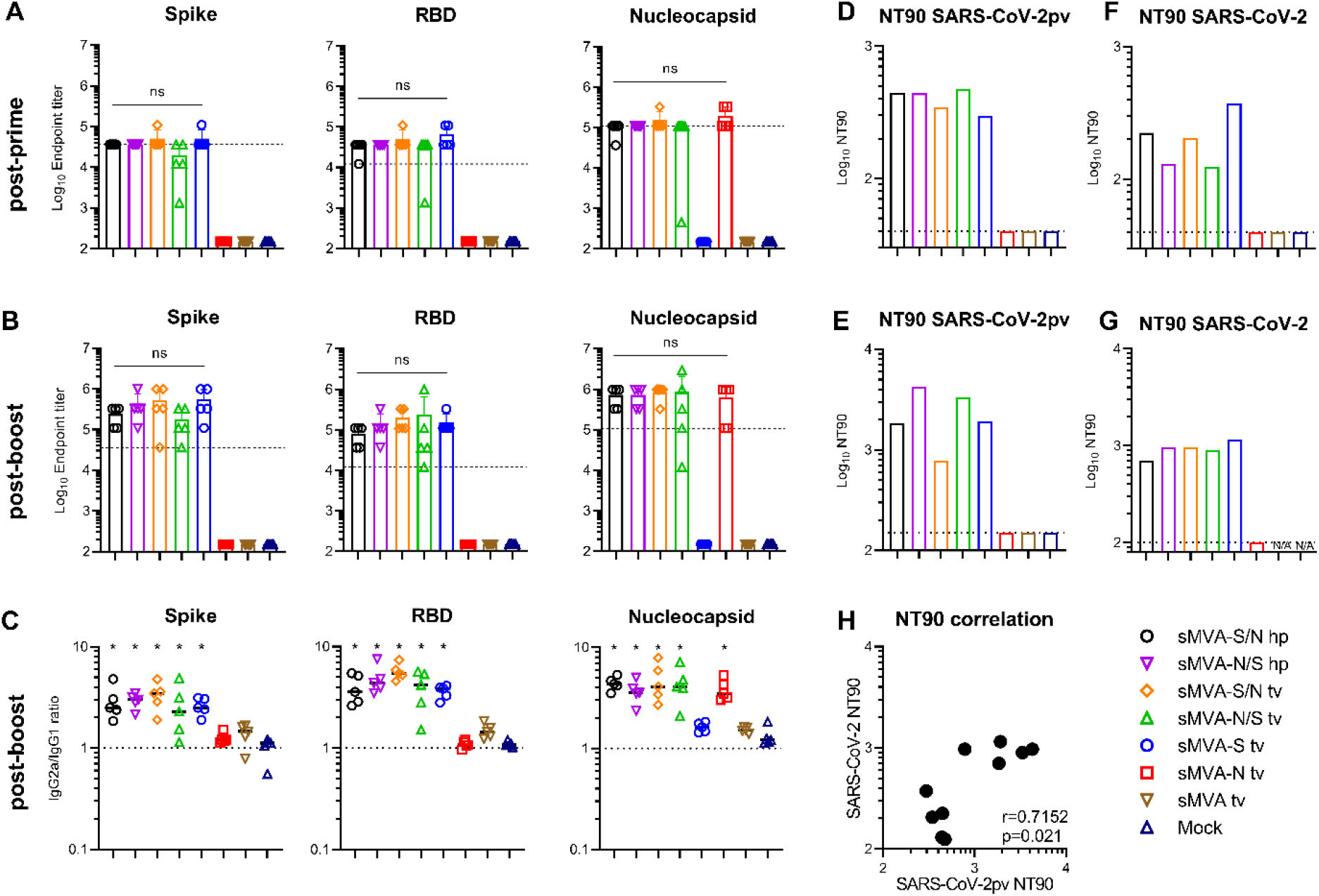
Humoral immune responses stimulated by sMVA-CoV2 vectors. Balb/c mice immunized twice in a three week interval with 5×10^7^ PFU of the single and double recombinant sMVA-CoV2 vectors derived with FPV HP1.441 (sMVA-S/N hp and sMVA-N/S hp) or TROVAC (sMVA-S/N tv, sMVA-N/S tv, sMVA-S tv, sMVA-N tv) were evaluated for SARS-CoV-2-specific humoral immune responses **A-B)** Binding antibodies. S, RBD, and N-specific binding antibodies induced by the vaccine vectors were evaluated after the first (A) and second (B) immunization by ELISA. Dashed lines in A and B indicate median binding antibody endpoint titers measured in convalescent human sera (Figure S4). One-way ANOVA with Tukey’s multiple comparison test was used to evaluate differences between binding antibody end-point titers. **C)** IgG2a/IgG1 isotype ratio. S-, RBD-, and N-specific binding antibodies of the IgG2a and IgG1 isotype were measured after the second immunization using 1:10,000 serum dilution, and absorbance reading was used to calculate IgG2a/IgG1 antibody ratio. One-way ANOVA with Dunnett’s multiple comparison test was used to compare each group mean IgG2a/IgG1 ratio to a ratio of 1 (balanced Th1/Th2 response). **D-G)** NAb responses. SARS-CoV-2-specific NAb (NT90 titer) induced by the vaccine vectors were measured after the first (D, F) and second (E, G) immunization against SARS-CoV-2 pseudovirus (pv) (D-E) or infectious SARS-CoV-2 virus (F-G) in pooled sera of immunized mice. Shown is the average NT90 measured in duplicate (D-E) or triplicate (F-G) infection. N/A=failed quality control of the samples. Dotted lines indicate lowest antibody dilution included in the analysis. **H)** SARS-CoV-2/SARS-CoV-2pv correlation analysis. Correlation analysis of NT90 measured in mouse sera after one and two immunizations using infectious SARS-CoV-2 virus and SARS-CoV-2pv. Pearson correlation coefficient (r) was calculated in H. *p<0.05. ns= not significant.

Potent SARS-CoV-2-specific NAb responses as assayed using pseudovirus were detected after the first immunization in all vaccine groups receiving the vectors encoding the S antigen (sMVA-S, sMVA-S/N, sMVA-N/S), and these NAb responses increased after the booster immunization (Figure 5D-E and S3D-E). Similar potent NAb responses as measured using pseudovirus were also observed in the vaccine groups using infectious SARS-CoV-2 virus (Figure 5F-G and S3F-G). We also evaluated the immune sera for potential antibody-dependent enhancement of infection (ADE) using THP-1 monocytes. These cells do not express the ACE2 receptor, but express Fc*γ* receptor II, which is considered the predominant mediator of ADE in SARS-CoV infection (*41*). THP-1 monocyte infection by SARS-CoV-2 pseudovirus was not promoted by the immune sera of any of the vaccine groups even at sub-neutralizing antibody concentrations (Figure S6), suggesting absence of Fc-mediated ADE by the vaccine-antibodies responses.

SARS-CoV-2-specific T cells evaluated after the second immunization by *ex vivo* antigen stimulation revealed both S- and N-specific T cell responses in the vaccine groups receiving the double recombinant vectors sMVA-S/N and sMVA-N/S. In contrast, mice receiving the single recombinant vectors sMVA-N or sMVA-S developed T cell responses only against either the N or S antigen (Figure 6A-D, Figures S7-8). High levels of cytokine-secreting (IFN*γ*, TNFα and IL-4) S-specific CD8+ T cells were measured in all vaccine groups immunized with the S-encoding sMVA-CoV2 vectors (Figure 6A). S-specific CD4^+^ T-cells mostly produced Th1 cytokines (IFNγ and TNFα), while production of Th2 cytokines (IL-4 and IL-10) did not increase following antigen stimulation (Figure 6C, S8), indicating a Th1-biased response. While activated N-specific CD8+ T cells were not detected at significant frequency (Figure 6B), N-specific IFN*γ* and to some degree TNFα-secreting CD4^+^ T cells were measured in all animals vaccinated with the single and double recombinant vectors encoding N (Figure 6D and S8). No significant differences were observed in the T cell levels of the single and double recombinant vaccine groups.

**Figure 6.**
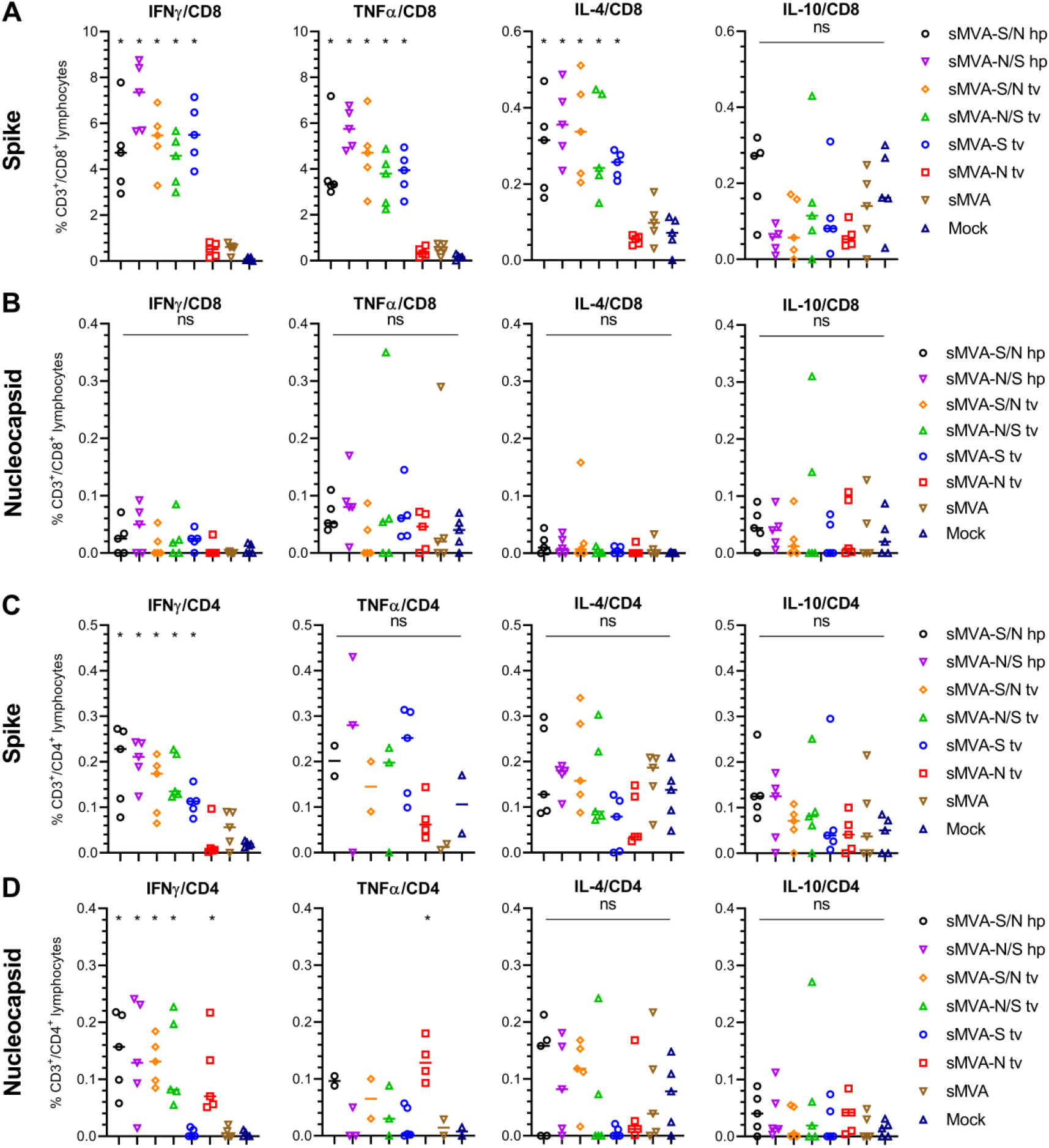
Cellular immune responses stimulated by sMVA-CoV2 vectors. Balb/c mice immunized twice in a three week interval with 5×10^7^ PFU of the single or double recombinant sMVA-CoV2 vectors derived with FPV HP1.441 (sMVA-S/N hp and sMVA-N/S hp) or TROVAC (sMVA-S/N tv, sMVA-N/S tv, sMVA-S tv, sMVA-N tv) were evaluated for SARS-CoV-2-specific cellular immune responses. Antigen-specific CD8+ (**A** and **B**) and CD4+ (**C** and **D**) T cell responses induced by the vaccine vectors after two immunizations were evaluated by flow cytometry for IFN*γ*, TNFα, IL-4 and IL-10 secretion following *ex vivo* antigen stimulation using SARS-CoV-2 S and N-specific peptide libraries. Due to technical issues, 1-3 animals/group were not included in the CD4/TNFα analysis in C and D. One-way ANOVA with Tukey’s multiple comparison test was used to compare differences in % of cytokine-specific T-cells between groups. *p<0.05. ns=not significant.

Stimulation of SARS-CoV-2-specific immune responses by both the S and N antigen was also evaluated in mice by co-immunization using the single recombinant vectors sMVA-S and sMVA-N at different doses. This study revealed similar SARS-CoV-2 antigen-specific humoral and cellular immune responses in vaccine groups receiving sMVA-S and sMVA-N alone or in combination (Figure S9-10). Altogether these results indicate that the sMVA-vectored S and N antigens when expressed alone or combined using a single vector or two separate vectors can stimulate potent SARS-CoV-2-specific humoral and cellular immune responses in mice.

## Discussion

We developed a novel vaccine platform based on a fully synthetic form of the highly attenuated and widely used MVA vector. In response to the ongoing global SARS-CoV-2 pandemic, we used this novel vaccine platform to rapidly produce sMVA vectors co-expressing SARS-CoV-2 S and N antigens and show that these vectors can induce potent SARS-CoV-2 antigen-specific humoral and cellular immune responses in mice, including potent NAb. These results highlight the feasibility to efficiently produce recombinant MVA vectors from chemically synthesized DNA and to rapidly develop a synthetic poxvirus-based vaccine candidate to prevent SARS-CoV-2 infection. We envision that this novel vaccine platform based on synthetic DNA will facilitate the development and clinical use of poxvirus vaccine vectors for infectious diseases and cancer.

Our strategy to produce a synthetic form of MVA using chemically synthesized DNA differs from the recently described approach to produce a synthetic horsepox virus vaccine vector (*42*). While our strategy to generate sMVA involves the use of three large circular DNA fragments (~60 kbp) with intrinsic HL and CR sequences (Figure 1), the approach by Noyce *et al.* to produce a synthetic horsepox vaccine involves the use of multiple smaller linear DNA fragments (~10-30 kbp) and the addition of terminal HL sequences (*42*). Because the three sMVA fragments can be used in a circular form for the sMVA reconstitution process they are easily maintained in *E. coli* as BACs and transferred to BHK cells for sMVA virus reconstitution without the need for additional purification steps or modifications. This feature greatly facilitates the insertion of heterologous antigen sequences into the sMVA DNA by highly efficient bacterial recombination techniques and to produce recombinant sMVA vaccine vectors. Additionally, the three-plasmid system provides the flexibility for rapid production of recombinant MVA harboring multiple antigens inserted into different MVA insertion sites, which can be particularly laborious when generating recombinant MVA by the conventional transfection/infection procedure (*3, 43*). Although the precise mechanism and order of events of the sMVA virus reconstitution using circular plasmids was not investigated, we demonstrate that the sMVA fragments efficiently recombine with one another and produce a synthetic form of MVA that is virtually identical to wtMVA in genome content, replication properties, host cell range, and immunogenicity.

In contrast to most other currently employed SARS-CoV-2 vaccine approaches that solely rely on the S antigen, our SARS-CoV-2 vaccine approach using sMVA employs immune stimulation by S and N antigens, which both are implicated in protective immunity (*20, 26*). The observation that the sMVA-CoV2 vectors co-expressing S and N antigens can stimulate potent NAb against SARS-CoV-2 pseudovirus and infectious virions suggests that they can elicit antibodies that are considered effective in preventing SARS-CoV-2 infection and disease (*16, 18, 20, 21*). We show that the vaccine vectors stimulate a Th1-biased antibody and cellular immune response, which is considered the preferred antiviral adaptive immune response to avoid vaccine-associated enhanced respiratory disease (*44, 45*). We did not find any evidence for Fc-mediated ADE promoted by the vaccine-induced immune sera, suggesting that antibody responses induced by the vaccine vectors bear minimal risk for ADE-mediated immunopathology, a general concern in SARS-CoV-2 vaccine development (*44, 45*). In addition, based on findings with other viruses associated with ADE, the stimulation of Th1 immunity with a strong T cell response component appears to be the way forward to develop an effective SARS-CoV-2 vaccine candidate (*46*).

Other immune responses besides NAb targeting the S antigen may contribute to protection against SARS-CoV-2 infection, which is highlighted by the finding that even patients without measurable NAb can recover from SARS-CoV-2 infection (*20*). While antibodies could be particular important to prevent initial SARS-CoV-2 acquisition, T cell responses may impose an additional countermeasure to control sporadic virus spread at local sites of viral infection, thereby limiting virus transmission. Our dual recombinant vaccine approach based on sMVA to induce robust humoral and cellular immune responses to S and N antigens may provide protection against SARS-CoV-2 infection beyond other vaccine approaches that solely employ the S antigen. Our results warrant further preclinical testing of a sMVA vaccine candidate for protective efficacy in animal models towards rapid advancement into phase 1 clinical testing.

## Materials and Methods

### Cells and Viruses

BHK-21 (CCL-10), A549 (CCL-185), HeLa (CCL-2), 293T (CRL-1573), 143B (CRL-8303), MRC-5 (CCL-171), HEK293/17 (CRL11268), THP-1 (TIB-202), ARPE-19 (CRL-2302) were purchased from the American Type Culture Collection (ATCC) and grown according to ATCC recommendations. CEF were purchased from Charles River (10100795) and grown in minimum essential medium (MEM) with 10% FBS (MEM10). HEK293T/ACE2 were a kind gift of Pamela J. Bjorkman (*47*). We acknowledge Bernard Moss (LVD, NIAID, NIH) for the gift of wtMVA (NIH Clone 1) that was used solely as a reference standard. To produce sMVA and wtMVA virus stocks, CEF were seeded in 30×150mm tissue culture dishes, grown to ~70-90% confluency, infected at 0.02 multiplicity of infection (MOI) with sMVA or wtMVA. Two days post infection, ultra-purified virus was prepared by 36% sucrose density ultracentrifugation and virus resuspension in 1 mM Tris-HCl (pH 9) (*48*). Virus stocks were stored at −80°C. Virus titers were determined on CEF by immunostaining of viral plaques at 16-24 h post infection using polyclonal Vaccinia antibody. FPV stocks were produced following propagation on CEF using FPV strain TROVAC (ATCC VR-2553) (*35*) or HP1.441 (*36*), kindly provided by Bernard Moss. FPV titers were evaluated on CEF by virus plaque determination.

### Construction of sMVA fragments

The three ~60 kbp sMVA fragments (F1-F3; Figure 1) comprising the complete MVA genome sequence reported by Antoine *et al.* (NCBI Accession# U94848) (*4*) were constructed as follows: sMVA F1 contained base pairs 191-59743 of the MVA genome sequence; sMVA F2 comprised base pairs 56744-119298 of the MVA sequence; and sMVA F3 included base pairs 116299-177898 of the reported MVA genome sequence (*4*). A CR/HL/CR sequence arrangement composed of 5’-TTT TTT TCT AGA CAC TAA ATA AAT A*GT AAG ATT AAA TTA ATT ATA AAA TTA TGT ATA TAA TAT TAA TTA TAA AAT TAT GTA TAT GAT TTA CTA ACT TTA GTT AGA TAA ATT AAT AAT ACA TAA ATT TTA GTA TAT TAA TAT TAT AAA TTA ATA ATA CAT AAA TTT TAG TAT ATT AAT ATT ATA TTT TAA A* TA TTT ATT TAG TGT CTA GAA AAA AA-3’ was added in the same orientation to both ends of each of the sMVA fragments, wherein the italicized letters indicate the duplex copy of the MVA terminal HL sequence and the underlined letters indicate the CR sequences. Notably, the CR/HL/CR sequences incorporated at the ITRs of sMVA F1 and F3 were added in identical arrangement as the CR/HL/CR sequences occur at the ITRs at the genomic junctions of putative MVA replication intermediates (*4*). The sMVA fragments were produced and assembled by Genscript using chemical synthesis, combined with a yeast recombination system. All sMVA fragments were cloned into a yeast shuttle vector, termed pCCI-Brick, which contains a mini-F replicon for stable propagation of large DNA fragments as low copy BACs in *E. coli.* sMVA F1 and F3 were cloned and maintained in EPI300 *E. coli* (Epicentre), while sMVA F1 was cloned and maintained in DH10B *E. coli* (Invitrogen).

### Antigen insertion

SARS-CoV-2 S and N antigen sequences were inserted into the sMVA fragments by *En passant* mutagenesis in GS1783 *E. coli* cells (*49, 50*). Briefly, transfer constructs were generated that consisted of the S or N antigen sequence with upstream mH5 promoter sequence and downstream Vaccinia transcription termination signal (TTTTTAT), and a kanamycin resistance cassette flanked by a 50 bp gene duplication was introduced into the antigen sequences. The transfer constructs were amplified by PCR with primers providing ~50 bp extensions for homologous recombination and the resulting PCR products were used to insert the transfer constructs into the sMVA DNA by a first Red-recombination reaction (*49, 50*). Primers 5’-AAA AAA TAT ATT ATT TTT ATG TTA TTT TGT TAA AAA TAA TCA TCG AAT ACG AAC TAG TAT AAA AAG GCG CGC C-3’ and 5’-GAA GAT ACC AAA ATA GTA AAG ATT TTG CTA TTC AGT GGA CTG GAT GAT TCA AAA ATT GAA AAT AAA TAC AAA GGT TC-3’ were used to insert the N antigen sequence into the Del2 site. Primers 5’-ATA TGA ATA TGA TTT CAG ATA CTA TAT TTG TTC CTG TAG ATA ATA ACT AAA AAT TTT TAT CTA GTA TAA AAA GGC GCG CC-3’ and 5’-GGA AAA TTT TTC ATC TCT AAA AAA AGA TGT GGT CAT TAG AGT TTG ATT TTT ATA AAA ATT GAA AAT AAA TAC AAA GGT TC-3’ were used to insert the S antigen sequence into the IGR69/70 insertion site primers. Primers 5’-TTG GGG AAA TAT GAA CCT GAC ATG ATT AAG ATT GCT CTT TCG GTG GCT GGT AAA AAA TTG AAA ATA AAT ACA AAG GTT C-3’ and 5’-ACA AAA TTA TGT ATT TTG TTC TAT CAA CTA CCT ATA AAA CTT TCC AAA TAC TAG TAT AAA AAG GCG CGC C-3’ were used to insert the S or N antigen sequence into the Del3 site. Underlined letters indicate the sequences used to produce ~50 bp extensions for homologous recombination. The S and N antigen sequences were based on the SARS-CoV-2 reference strain (NCBI Accession# NC_045512) and codon-optimized for Vaccinia (*10, 38*). Inserted antigen sequences were verified by PCR, restriction enzyme digestion, and sequencing.

### sMVA virus reconstitution

sMVA virus reconstitution from the three sMVA DNA plasmids in BHK cells using FPV as a helper virus was performed as follows (*8–10*). The three sMVA DNA plasmids were isolated from *E. coli* by alkaline lysis (*51*) and co-transfected into 60-70% confluent BHK cells grown in 6-well plate tissue culture plates using Fugene HD transfection reagent (Roche) according to the manufacturer’s instructions. At 4 hours post transfection, the cells were infected with approximately 0.1-1 MOI of FPV to initiate the sMVA virus reconstitution. The transfected/infected BHK cells were grown for 2 days and then every other day transferred, re-seeded, and grown for additional two days in larger tissue culture formats over a period of 8-12 days until most or all of the cells showed signs of sMVA virus infection. Using this procedure, characteristic MVA viral plaque formation and cytopathic effects (CPEs) indicating sMVA virus reconstitution was usually detected at 4-8 days post transfection/infection. Fully infected BHK cell monolayers were usually visible at 8-12 days post transfection/infection. sMVA virus from infected BHK cell monolayers was prepared by conventional freeze/thaw method and passaged once on BHK cells before producing ultra-purified virus stocks on CEF. sMVA or recombinant sMVA-CoV-2 vectors were reconstituted either with FPV HP1.441 (sMVA hp, sMVA-N/S, sMVA-S/N hp) or TROVAC (sMVA tv1 and tv2, sMVA-S tv, sMVA-N tv, sMVA-N/S tv, sMVA-S/N tv).

### Host cell range

sMVA and wtMVA host cell range using various human cell lines (HeLa, 293T, MRC-5, A549, and 143B) BHK cells, and CEF was determined as follows. The cells were seeded in 6-well plate tissue culture format and at 70-90% confluency infected in duplicates with 0.01 MOI of sMVA or wtMVA using MEM2. At 2 hours post infection, the cells were washed twice with PBS and incubated for two days in normal growth medium (as described under cells and viruses). After the incubation period, virus was prepared by conventional freeze/thaw method and the virus titers of each duplicate infection was determined in duplicate on CEF.

### Replication kinetics

To compare the replication kinetics of sMVA and wtMVA, CEF or BHK cells were seeded in 6 well-plate tissue culture format and at 70-90% confluency infected in triplicates at 0.02 MOI with sMVA or wtMVA using MEM2. After 2 hours of incubation, the cells were grown in MEM10. At 24 and 48 hours post infection, virus was prepared by freeze/thaw method and the virus titers of each triplicate infection and the inoculum was determined in duplicate on CEF.

### Plaque size analysis

To compare the plaque size of sMVA virus and wtMVA, CEF or BHK cells were seeded in 6-well plate tissue culture format and at 70-90% confluency infected with 0.002 MOI with sMVA or wtMVA using MEM2. After 2 hours of incubation, MEM10 was added and the cells were grown for 16-24 hours. The cell monolayers were stained with Vaccinia virus polyclonal antibody and viral plaques were imaged using Leica DMi8 inverted microscope and measured using LAS X software. The size of 25 viral plaques per sMVA or wtMVA was calculated using the formula *Area*= *π*×*a*×*b*, where *a* and *b* are the major and minor radius of the ellipse, respectively. PCR analysis

To characterize the viral DNA of the sMVA vectors by PCR, CEF were seeded in 6-well plate tissue culture format and at 70-90% confluency infected at 5 MOI with sMVA or wtMVA. DNA was extracted at 16-24 hours post infection by the DNA Easy Blood and Tissue Kit (Qiagen) according to the manufacturer’s instructions. All PCR reactions were performed with Phusion polymerase (ThermoFischer Scientific). Primers 5’-TCG TGG TGT GCC TGA ATC G-3’ and 5’-AGG TAG CGA CTT CAG GTT TCT T-3’ were used to detect MVA ITR sequences; primers 5’-TAT CCA CCA ATC CGA GAC CA-3’ and 5’-CCT CTG GAC CGC ATA ATC TG-3’ were used to verify the transition from the left ITR into the unique region; primers 5’-AGG TTT GAT CGT TGT CAT TTC TCC-3’ and 5’-AGA GGG ATA TTA AGT CGA TAG CCG-3’ were used to verify the Del2 site with or without inserted N antigen sequence; primers 5’-TGG AAT GCG TTC CTT GTG C-3’ and 5’-CGT TTT TCC CAT TCG ATA CAG-3’ with binding sites flanking the F1/F2 homologous sequences were used to verify the F1/F2 recombination site; primers 5’-TAT AGT CTT TGT GGC ATC CGT TG-3’ and 5’-ACC CAA ACT TTA GTA AGG CCA TG-3’ were used to verify the IGR69/70 insertion site with or without inserted S antigen; primers 5’-ATA AGC GTT GTC AAA GCG GG-3’ and 5’-AGG AAA TAG AAA TTG TTG GTG CG-3’ with binding sites flanking the F2/F3 homologous sequences were used to verify the F2/F3 recombination site; primers 5’-ACA TTG GCG GAC AAT CTA AAA AC-3’ and 5’-ATC ATC GGT GGT TGA TTT AGT AGT G-3’ were used to verify the Del3 insertion site with and without inserted S or N antigen sequences; primers 5’-TAT CCA CCA ATC CGA GAC CA-3’ and 5’-GTC TGT CCG TCT TCT CTA TTG TTT A-3’ were used to verify the transition from the unique region into the right ITR; and primers 5’-TTA ACT CAG TTT CAA TAC GGT GCA G-3 and 5’-TGG GGT TTC TTC TCA GGC TAT C-3’ were used to detect the *SopA* element of the BAC vector. Restriction pattern analysis

BHK cells were seeded in 20×150 mm tissue culture dishes, grown to ~70-90% confluency, and infected at 0.01 MOI with wtMVA, sMVA tv1, or sMVA tv2. Ultra-purified virus was prepared two days post-infection as previously described (*48*). Viral DNA (vDNA) was phenol/chloroform extracted, followed by ethanol precipitation as previously described(*52*). DNA concentration and A_260_/A_280_ ratios were determined using NanoVue (GE Healthcare Bio-sciences Corp). 10 μg of vDNA were digested with 3 units of either KpnI or XhoI, followed by visualization on 0.5% EtBr-stained agarose gel that was run at 2.4v/cm, overnight.

### Sequencing of sMVA fragments and genome

PacBio Long Read Sequencing analysis confirmed the integrity of the sMVA fragments and sMVA genome, including a single point mutation in a non-coding determining region at 3 base pairs downstream of 021L (*4*) that was found both in sMVA F1 and in reconstituted sMVA. Briefly, 5 ug of fragmented DNAs were converted to SMRTbell libraries using the SMRTbell Template Prep Kit 1.0 (PacBio). The libraries were size-selected (7-kb size cutoff) with BluePippin (Sage Science). The size-selected libraries were loaded to SMRT cells using MagBeads loading and sequenced on a PacBio RSII with 10 hour movie. Read demultiplexing, mapping to the reference sequence (Vaccinia virus strain Ankara U94848.1), and variants calling were performed using the SMRT Link (v6.0.0.47841). The identified variants were visually inspected in SMRT view Genome Browser for confirmation. De novo assembly was done using either canu v1.7.1 or wtdbg2 v2.5. The 5’ start position of the assembled contig was edited by comparing to the U94848.1 reference.

### Immunblot analysis

BHK cells infected at 5 MOI were harvested 24-post infection. Proteins were solubilized in PBS with 0.1% Triton X-100, supplemented with protease inhibitor, then reduced and denatured in Laemmli buffer containing DTT and boiled at 95°C for ~10 minutes. Proteins were resolved on a 4-20% Mini Protean TGX gradient gel (BioRad), and transferred onto PVDF membrane. S protein was probed with anti-SARS-CoV-1 S1 subunit rabbit polyclonal antibody (40150-T62-COV2, Sino Biological); N protein was probed with anti-SARS-CoV1 NP rabbit polyclonal antibody (40413-T62, Sino Biological). Vaccinia BR5 protein was probed as a loading control. Anti-rabbit polyclonal antibody conjugated with horseradish peroxidase (Sigma-Aldrich) was used as a secondary antibody and protein bands were visualized with chemiluminescent substrate (ThermoFisher).

### Flow cytometry

HeLa cells were seeded in a 6-well plate (5×10^5^/well) and infected the following day with sMVA vaccine candidates at an MOI of 5. Following an incubation of 6 hours, cells were detached with non-enzymatic cell dissociation buffer (13151014, GIBCO). Cells were either incubated directly with primary antibody or fixed and permeabilized prior to antibody addition. Anti-SARS-CoV-1 S1 mouse (40150-R007, Sino Biological) and S2 rabbit (GTX632604, GeneTex) monoclonal antibodies, anti-SARS-CoV-1 N rabbit monoclonal antibody (40143-R001, Sino Biological), and anti-vaccinia rabbit polyclonal antibody (9503-2057, Bio Rad) were used in dilution 1:2,000. One hour later anti-mouse or anti-rabbit Alexa Fluor 488-conjugated secondary antibodies (A11001, A21206; Invitrogen) were added to the cells at a dilution of 1:4,000. Live cells were ultimately fixed with 1% paraformaldehyde (PFA).

### Immunofluorescence

BHK or HeLa cells were grown on glass coverslips and infected with sMVA or recombinant sMVAs encoding S and/or N proteins at an MOI of 5 for 6 hours at 37°C in a humidified incubator (5% CO_2_). After infection, cells were fixed for 1 m5in in 2% PFA and then directly permeabilized by addition of ice cold 1:1 acetone/methanol for 5 min on ice. Cells were blocked for 1 hr with 3% BSA at room temperature, incubated with primary antibody mix (1:500) against the S2 subunit or N for 1 hr at 37°C, and then incubated with Alexa-conjugated secondary antibodies (ThermoFischer) (1:2000) for 1 hr at 37°C, with washing (PBS + 0.1% Tween20) between each step. For detection of cell membranes and nuclei, cells were incubated with Alexa-conjugated wheat germ agglutinin at 5 μg/mL (Thermo Fisher) and DAPI for 10 minutes at room temperature. Coverslips were washed and mounted onto slides with Fluoromount-G (SouthernBiotech). Microscopic analysis was performed using a laser-scanning confocal microscope (Zeiss, LSM700). Images were acquired and processed using Zen software (Zeiss).

### Mouse immunization

The Institutional Animal Care and Use Committee (IACUC) of the Beckman Research Institute of City of Hope (COH) approved protocol 20013 assigned for this study. All study procedures were carried out in strict accordance with the recommendations in the Guide for the Care and Use of Laboratory Animals and the Public Health Service Policy on the Humane Care and Use of Laboratory Animals. 6 weeks old C57BL/6 (C57BL/6J, 000664) or Balb/c (BALB/cJ, 000651) were purchased from the Jackson laboratories. C57BL/6 Nramp were bred at the City of Hope animal facility. Mice (N=4-5) were immunized twice in three weeks interval by intraperitoneal route with 5×10^7^ PFU (high dose) or 1×10^7^ PFU (low dose) of sMVA, wtMVA, or sMVA-CoV2 vectors. To determine immune stimulation by both the S and N antigen when using separate vectors (Figures S9-10), mice were co-immunized via the same immunization schedule and route with half of the high (2.5×10^7^ PFU) or low dose (0.5×10^7^ PFU) of each of the vaccine vectors. Blood samples for humoral immune analysis were collected by retro-orbital bleeding two weeks post-prime and one-week post booster immunization. Splenocytes for cellular immune analysis were collected at one week post booster immunization and were isolated by standard procedure after animals were humanely euthanized.

### Binding antibodies

Binding antibodies in mice immunized with sMVA, wtMVA, or sMVA-CoV2 vectors were evaluated by ELISA. ELISA plates (3361, Corning) were coated overnight with 1 μg/ml of MVA expressing Venus fluorescent marker (*9*), S (S1+S2, 40589-V08B1, Sino Biological), RBD (40592-V08H, Sino Biological) or N (40588-V08B, Sino Biological). Plates were blocked with 3% BSA in PBS for 2 hours. Serial dilutions of the mouse sera were prepared in PBS and added to the plates for two hours. After washing the plate, 1:3,000 dilution of HRP-conjugated anti-mouse IgG secondary antibody (W402B, Promega) was added and incubated for one additional hour. Plates were developed using 1-Step Ultra TMB-ELISA (34028, Thermo Fisher) for one to two minutes after which the reaction was stopped with 1M H_2_SO_4_. Plates were read at 450 nanometers wave length using FilterMax F3 microplate reader (Molecular Devices). Binding antibodies endpoint titers were calculated as the latest serum dilution to have an absorbance higher than 0.1 absorbance units (OD) or higher than the average OD in mock immunized mice plus 5 times the standard deviation of the OD in the same group at the same dilution. For evaluation of the IgG2a/IgG1 ratio, mouse sera were diluted 1:10,000 in PBS. The assay was performed as described above except for the secondary antibodies (1:2,000. goat Anti-Mouse IgG2a cross absorbed HRP antibody, Southern biotech, 1083-05; Goat anti-Mouse IgG1 cross absorbed HRP antibody, Thermo Fisher, A10551). The IgG2a/IgG1 ratio was calculated by dividing the absorbance read in the well incubated with the IgG2a secondary antibody divided the absorbance for the same sample incubated with the IgG1 antibody.

### MVA neutralization assay

ARPE-19 cells were seeded in 96 well plates (1.5×10^4^ cells/well). The following day, serial dilutions of mouse sera were incubated for 2 hours with MVA expressing the fluorescent marker Venus (*9*) (1.5×10^4^ PFU/well). The serum-virus mixture was added to the cells in duplicate wells and incubated for 24 hours. After the 24 hours incubation period, the cells were imaged using Leica DMi8 inverted microscope. Pictures from each well were processed using Image-Pro Premier (Media Cybernetics) and the fluorescent area corresponding to the area covered by MVA-Venus infected cells was calculated.

### SARS-CoV-2 pseudovirus production

The day before transfection, HEK293T/17 were seeded in a 15 cm dish at a density of 5×10^6^ cells in DMEM supplemented with 10% heat inactivated FBS, non-essential amino acids, HEPES, and glutamine (*53*). Next day, cells were transfected with a mix of packaging vector (pALDI-Lenti System, Aldevron), luciferase reporter vector and a plasmid encoding for the wild type SARS-CoV2 Spike protein (Sino Biological) or vesicular stomatitis virus G (VSV-G, Aldevron), using FuGENE6 (Roche) as a transfection reagent : DNA ratio of 3:1, according to manufacturer’s protocol. Sixteen hours post-transfection, the media was replaced and cells were incubated for an additional 24-72 hours. Supernatants were harvested at 24-, 48- and 72 hours, clarified by centrifugation at 1,500 RPM for 5 minutes and filtered using a sterile 0.22 μm pore size filter. Clarified lentiviral particles were concentrated by ultracentrifugation at 20,000 RPM for 2 hours at 4°C. The pellet was resuspended in DMEM containing 2% heat inactivated-FBS and stored overnight at 4°C to allow the pellet to completely dissolve. Next day, samples were aliquoted, snap frozen and stored at −80°C for downstream assays.

### SARS-CoV-2 pseudotype neutralization and ADE assay

Levels of p24 antigen in the purified SARS-CoV-2 pseudotype solution was measured by ELISA (Takara). Mouse sera were heat inactivated, pooled and diluted at a linear range of 1:100 to 1:50,000 in complete DMEM. For the neutralization assay, diluted serum samples were pre-incubated overnight at 4°C with SARS-CoV-2-Spike pseudotyped luciferase lentiviral vector, normalized to 100 ng/mL of p24 antigen. HEK293T cells overexpressing ACE-2 receptor were seeded the day before transduction at a density of 2×10^5^ cells per well in a 96-well plate in complete DMEM (*47*). Before infection, 5 μg/mL of polybrene was added to each well. Neutralized serum samples were then added to the wells and the cells were incubated for an additional 48 hours at 37°C and 5% CO_2_ atmosphere. Following incubation, cells were lysed using 40 μL of Luciferase Cell Culture Lysis 5x Reagent per well (Promega). Luciferase activity was quantified using 100 μL of Luciferase Assay Reagent (Promega) as a substrate. Relative luciferase units (RLU) were measured using a microplate reader (SpectraMax L, Molecular Devices) at a 570 nm wave length. The percent neutralization titer for each dilution was calculated as follows: NT = [1-(mean luminescence with immune sera/mean luminescence without immune sera)] x 100. The titers that gave 90% neutralization (NT90) were calculated by determining the linear slope of the graph plotting NT versus serum dilution by using the next higher and lower NT. In all the experiments RLU of uninfected cells was measured and was always between 50 and 90.

For the ADE assay, THP1 cells were seeded at a confluency of 2×10^6^ cells/mL in a 96 well plate and co-incubated for 48 hours with serum samples diluted at 1:5,000 or 1:50,000 in the presence of SARS-CoV-2-Spike pseudotyped or VSV-G luciferase lentiviral vector, normalized to 100 ng/mL of p24 antigen. Following incubation, cells were lysed using 100 μL of ONE-Glo Luciferase Assay System per well (Promega). RLU were measured as above.

### SARS-CoV-2 focus reduction neutralization test (FRNT)

FRNT assay was performed as described recently (*54*). Briefly, HeLa-ACE2 cells were seeded in 12 μL complete DMEM at a density of 2×10^3^ cells per well. In a dilution plate, pooled mouse serum was diluted in series with a final volume of 12.5 μL. Then 12.5 μL of SARS-CoV-2 was added to the dilution plate at a concentration of 1.2×10^4^ pfu/mL.

After 1 h incubation, the media remaining on the 384-well plate was removed and 25 μL of the virus/serum mixture was added to the 384-well plate. The plate was incubated for 20 h after which the plate was fixed for 1h. Each well was then washed three times with 100 μL of 1xPBS 0.05% tween. 12.5 μL of human polyclonal sera diluted 1:500 in Perm/Wash buffer (BD Biosciences 554723) were added to each well in the plate and incubated at RT for 2 h. Each well was further washed three times and peroxidase goat anti-human Fab (Jackson Scientific) was diluted 1:200 in Perm/Wash buffer, then added to the plate and incubated at RT for 2 h. The plate was then washed three times and 12.5 μL of Perm/Wash buffer was added to the plate then incubated at RT for 5 min. The Perm/Wash buffer was removed and TrueBlue peroxidase substrate was immediately added (Sera Care 5510-0030). Sera were tested in triplicate wells. Normal human plasma was used as negative controls for serum screening.

### SARS-CoV-2 convalescent plasma samples

IBC Protocol 20004 approved the use of SARS-CoV-2 convalescent plasma. Anonymized plasma samples of SARS-CoV-2 convalescent individuals (N=19) were obtained from UCSD. Individuals were confirmed to be infected in the previous three to ten weeks by PCR and lateral flow assay. All individuals were symptomatic with mild to moderate-severe symptoms. Serum samples (DS-626-G and DS-626-N, Seracare) purchased before SARS-CoV-2 pandemic were used as a negative control. SARS-CoV-2-specific binding antibodies in plasma samples were measured as described above. Cross-adsorbed goat anti-human IgG (H+L) secondary antibody (A18811, Invitrogen) was used at a dilution of 1:3,000.

### T cell analysis

Spleens were harvested and dissociated using a cell mesh following which blood cells were removed using RBC Lysis Buffer (BioLegend). 2.5×10^6^ splenocytes were stimulated with S or N peptide libraries (GenScript, 15mers with 11aa overlap, 1μg/ml), 0.1% DMSO, or phorbol myristate acetate (PMA)-ionomycin (BD Biosciences) for 1.5 h at 37°C. Anti-mouse CD28 and CD49d antibodies (1μg/ml; BioLegend) were added as co-stimulation. Brefeldin A (3μg/ml; eBioscience) was added, and the cells were incubated for additional 16 h at 37°C. Cells were fixed using Cytofix buffer (BD Biosciences) and surface staining was performed using fluorescein isothiocyanate (FITC)-conjugated anti-mouse CD3 (Clone 17A2, 555274, BD), BV650 anti-mouse CD8a (Clone 53-6.7, 563234, BD). Following cell permeabilization using Cytoperm buffer (BD Biosciences), ICS was performed using allophycocyanin (APC)-conjugated anti-mouse IFN-γ (Clone XMG1.2, 554413, BD), phycoerythrin (PE)-conjugated anti-mouse TNF-α (Clone MP6-XT22, 554419, BD), and PE-CF594 anti-mouse IL-2 (BD Biosciences (Clone JES6-5H4, 562483, BD). In experiments testing double recombinants SARS-CoV2 vectors IL-2 antibody was not included and PE-CF594 anti-mouse IL-4 (clone 11B11, 562450, BD) and BV421 rat anti mouse IL-10 (clone JES5-16E3, 563276, BD) were added. Events were acquired using a BD FACSCelesta flow cytometer (2×10^5^ cells/tube). Analysis was performed using FlowJo. Antigen specific T cells were identified by gating on size (FSC vs SSC), doublet negative (FSC-H vs FSC-A), CD3+, CD8+/CD4+. Cytokine positive responses are presented after subtraction of the background response detected in the corresponding unstimulated sample (media added with Brefeldin A one hour after beginning of mock stimulation) of each individual mouse sample.

### Cytokines ELISA

Splenocytes (1×10^6^) from immunized mice were incubated in v-bottom wells in the presence of 2μg/ml S or N peptide pools, or without stimulus in a volume of 200μl. 48 hours later, plates were centrifuged 2000 RPM for 10 minutes and cell supernatant was collected and stored at −80°C. Mouse TNF-alpha (MTA00B), Quantikine ELISA kit (R&D systems) was used according to manufacturer’s recommendations.

### Statistics

Statistical evaluation was pursued using GraphPad Prism (v8.3.0). For evaluation of differences in sMVA and wtMVA plaque area in BHK-21 and CEF cells and differences in sMVA and wtMVA host cell range, one-way ANOVA followed by Tukey’s and Dunnet’s multiple comparison tests were used, respectively. For sMVA and wtMVA growth kinetic analysis, mixed-effects model with the Geisser-Greenhouse correction, followed by Tukey’s multiple comparisons test were applied. For ELISAs, one-way ANOVA and Tukey’s multiple comparison tests were used to calculate differences in endpoint titers and group means between groups. For IgG2a/IgG1 ratio analysis, one-way ANOVA with Dunnett’s multiple comparison test was used to compare the IgG2a/IgG1 ratio measured in each group to a ratio of 1. Pearson correlation analysis was performed to calculate the correlation coefficient r and its significance. For T cell responses analysis, one-way ANOVA followed by Dunnett’s multiple comparisons test with a single pooled variance was used to compare the mean of each group.

## Acknowledgment

We thank Dr. Ildiko Csiki for her support and suggestions. We thank Dr. Ferdynand Kos and Weimin Tsai for their assistance with flow cytometry. We are grateful to Dr. Sara Gianella Weibel and Dr. Stephen Rawlings (UCSD) for contributing anonymized plasma from convalescent SARS-CoV-2 patients. We acknowledge Dr. Bernard Moss and the Laboratory of Viral Diseases of the NIAID for providing the FPV HP1.441 and plaque-purified MVA as a reference standard. We thank James Ricketts and Nathan Beutler (UCSD and Scripps Research) for their assistance with the SARS-CoV-2 FRNT assay. Special thanks go to Gloria Beauchamp and Shannon Dempsey for helping with the laboratory logistics.

