## Supplemental Figures for "Development of a Synthetic Poxvirus-Based SARS-CoV-2 Vaccine"

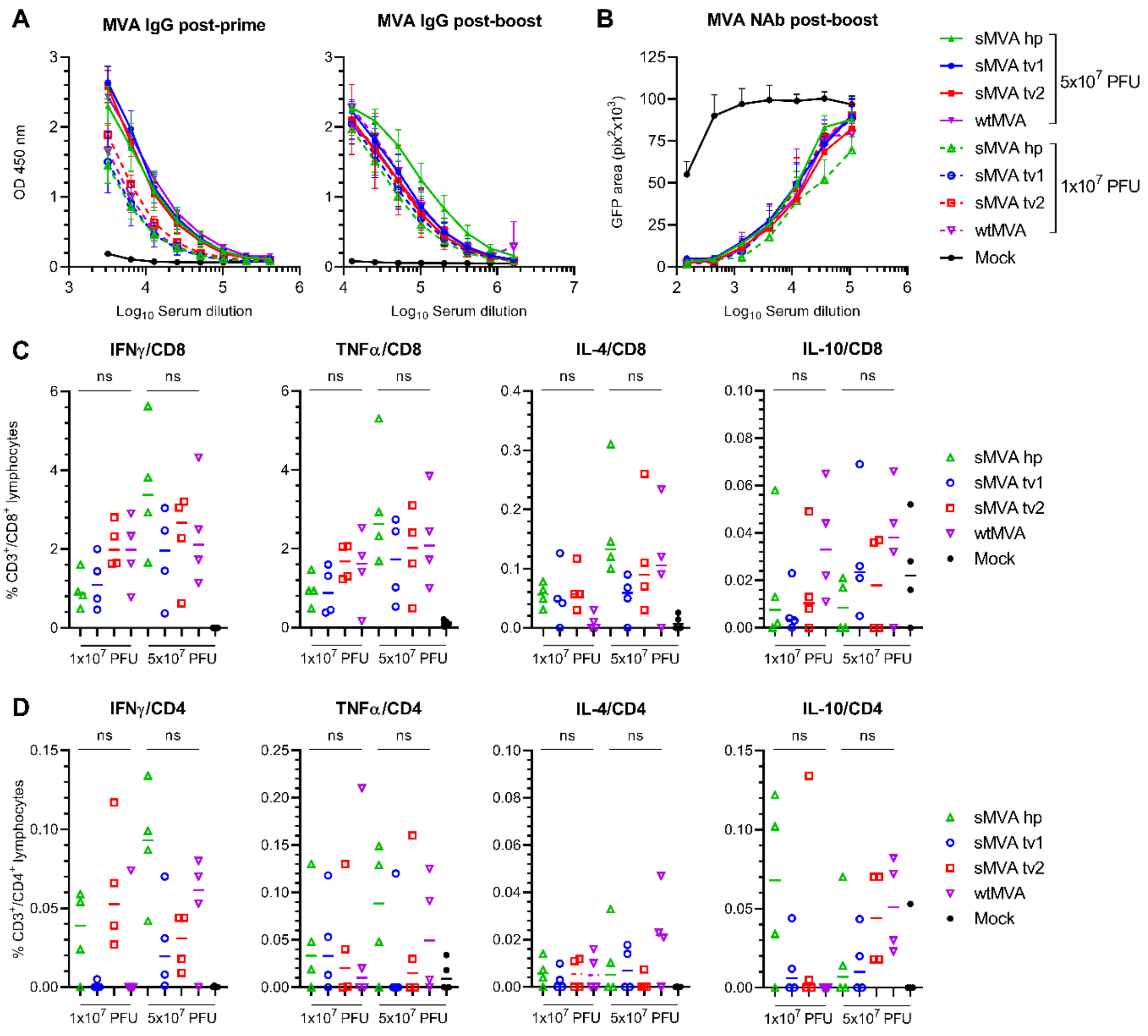


**Figure S1. Relative to Figure 3. sMVA immunogenicity *in vivo*.** sMVA derived either with FPV strain HP1.441 (sMVA hp) or with FPV strain TROVAC from two independent virus reconstitution (sMVA tv1 and sMVA tv2) was compared by *in vitro* analysis with wtMVA. C57BL/6 mice (N=4) were immunized twice in a three week interval with low ($1x10^7$ PFU) or high ($5x10^7$ PFU) dose of sMVA or wtMVA. Mock-immunized mice were used as controls **A)** Binding antibodies. Shown is the absorbance at 450 nm at different serum dilutions of MVA-specific binding antibodies (IgG titer) measured by ELISA after the first and second immunization in mice receiving sMVA or wtMVA. **B)** NAb responses. MVA-specific NAb titers induced by sMVA or wtMVA were measured after the booster immunization against wtMVA expressing a GFP marker. Shown is the measured GFP area of infected cells in square pixels ($pix^2x10^3$) at different serum dilutions **C-D)** T cell responses. MVA-specific CD8+ (C) and CD4+ (D) T cells expressing IFNγ, TNFα, IL-4, and IL-10 were measured after two immunizations with sMVA or wtMVA by flow cytometry following *ex vivo* antigen stimulation using Vaccinia A19L immunodominant peptides. Differences between groups were evaluated using one-way ANOVA with Tukey's multiple comparison test. ns = not significant.

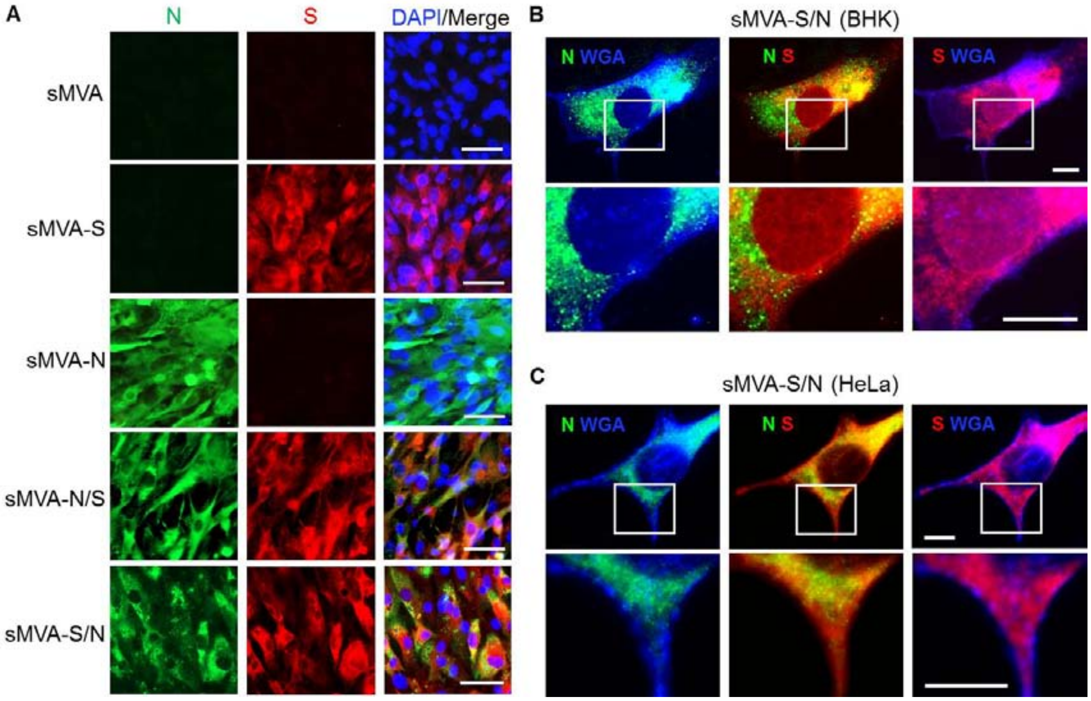


**Figure S2. Related to Figure 3. *In vitro* characterization of sMVA-CoV2 vectors.** S and N antigen expression by the single (sMVA-S and sMVA-N) and double (sMVA-S/N and sMVA-N/S) recombinant vaccine sMVA-CoV2 vectors – all derived with FPV HP1.441 – was evaluated in BHK (**A** and **B**) or HeLa (**C**) cells by immunofluorescent confocal imaging using N and S-specific antibodies. Fluorescently-conjugated wheat germ agglutinin (WGA) was used in B and C to stain the cell membrane. Magnified insets are found below images. Scale bars in A, 50 μm. Scale bars in B and C, 10 μm. All images represent two independent experiments with similar results.

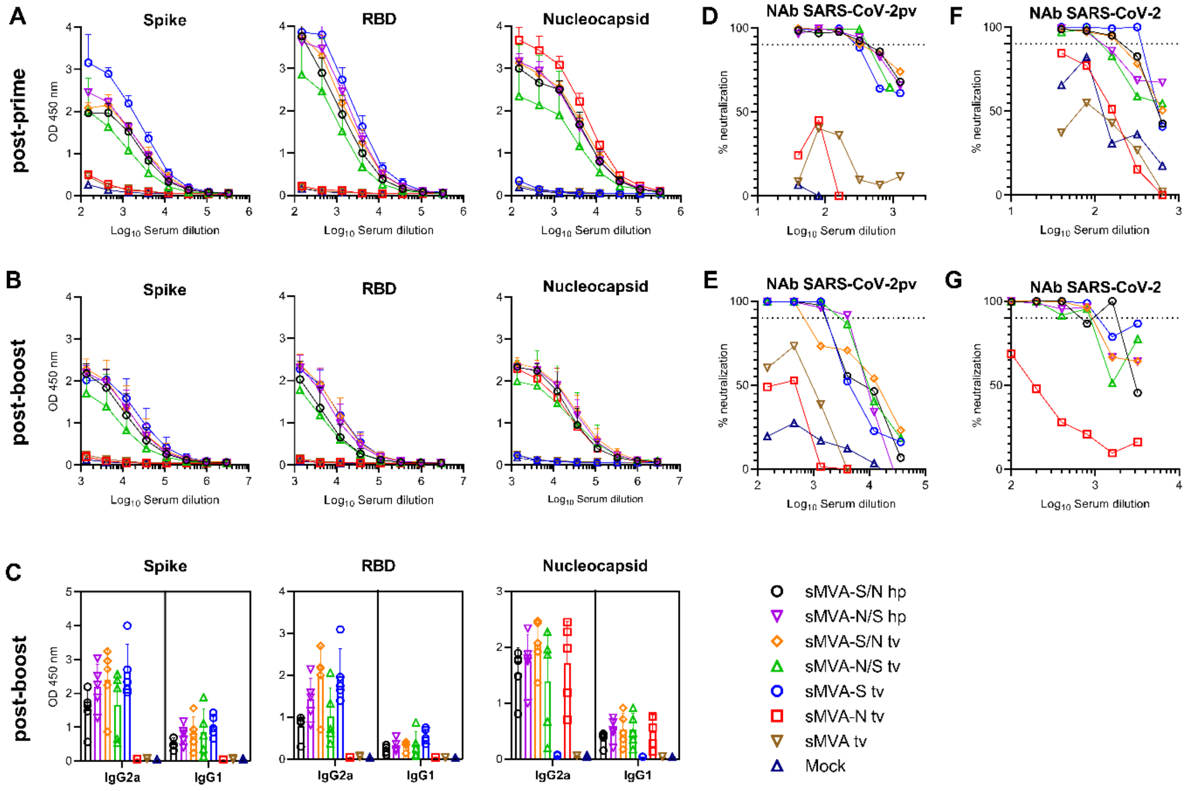


**Figure S3. Relative to Figure 5. Humoral immune responses induced by the sMVA-CoV2 vectors.** Shown are the antibody measurements in Balb/c mice (N=5) immunized twice in a three week interval with $5x10^7$ PFU of the single or double recombinant sMVA-CoV2 vectors derived with FPV HP1.441 (sMVA-S/N hp and sMVA-N/S hp) or TROVAC (sMVA-S/N tv, sMVA-N/S tv, sMVA-S tv, sMVA-N tv). **A-B)** Binding antibodies. Shown are S-, RBD-, and N-specific ELISA measurements at 450 nm using serial dilutions of serum collected two weeks post-prime (A) or one-week post-boost (B). **C)** IgG2a/IgG1 isotype ratio. Binding antibodies of the IgG2a and IgG1 isotypes were measured in serum of mice post-boost using a dilution of 1:10,000. **D-G)** NAb responses. Shown is the percent (%) of SARS-CoV-2pv (D-E) and infectious SARS-CoV-2 (F-G) neutralization measured in sera pooled from each group of immunized mice. Shown is the average % neutralization in duplicate (D-E) or triplicate (F-G) infection measured at different serum dilutions. Vaccine groups immunized with sMVA tv and PBS (mock) were not included in the analysis shown in G because of failure of quality control. Dotted lines mark 90% neutralization that was used to calculate NT90 in Figure 5.

.

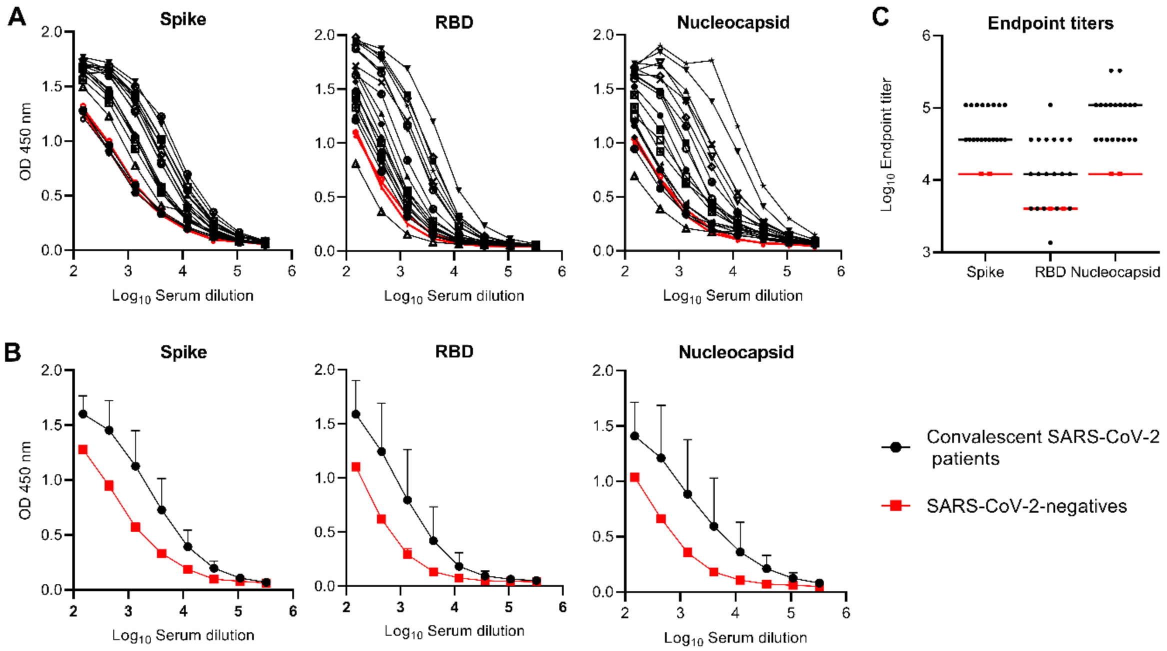


**Figure S4. Related to Figure 5. SARS-CoV-2-specific humoral immune responses in convalescent immune sera.** S-, RBD, and N-specific binding antibodies were measured via ELISA using serial dilutions of plasma samples from SARS-CoV-2 convalescent individuals. **A)** Binding antibody curves from individual samples (N=19). **B)** SARS-CoV-2 convalescent plasma binding curves were grouped together and compared to binding measured in samples (N=2) from SARS-CoV-2 negative individuals. **C)** Endpoint binding antibody titers to S, RBD, and N were calculated in individual plasma samples. Lines represent the median endpoint titers. Due to the limited number of SARS-CoV-2-negative samples evaluated, statistical analysis was not performed.

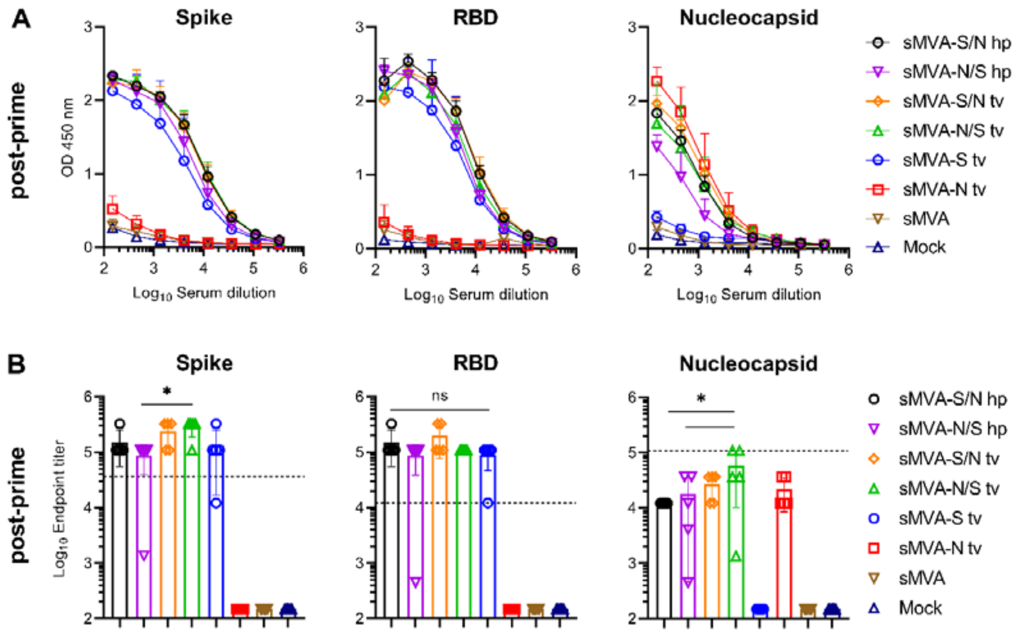


**Figure S5. Relative to Figure 5. Humoral immune responses induced by sMVA-CoV2 vectors.** C57BL/6 Nramp1 mice (N=5) were immunized with $5 \times 10^7$ PFU of the single and double recombinant sMVA-CoV2 vectors derived with FPV HP1.441 (sMVA-S/N hp and sMVA-N/S hp) or TROVAC (sMVA-S/N tv, sMVA-N/S tv, sMVA-S tv, sMVA-N tv) and evaluated for SARS-CoV-2-specific humoral immune responses **A-B)** Binding antibodies. S, RBD, and N-specific binding antibodies induced by the vaccine vectors were evaluated after the first immunization by ELISA. Dashed lines in B indicate median binding antibody endpoint titers that were measured in convalescent human sera (Figure S4). One-way ANOVA with Tukey's multiple comparison test was used to compare differences between binding antibody end-point titers in mice immunized with different vaccine vectors. *p<0.05. ns=not significant.

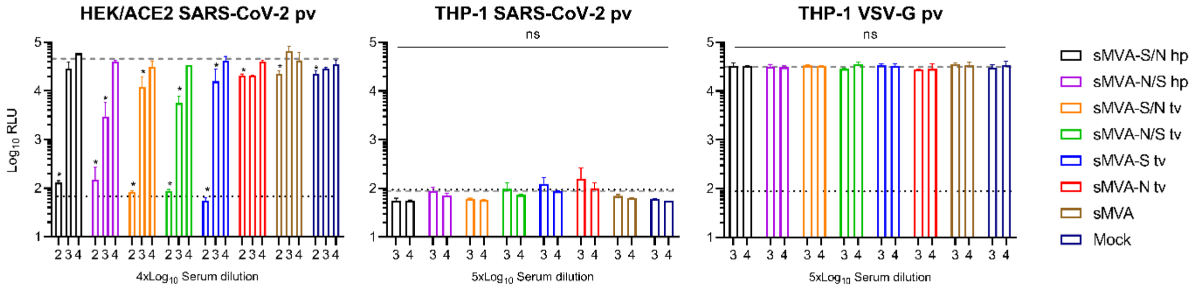


**Figure S6. Related to Figure 5. ADE assay.** The immune sera of Balb/c mice immunized with $5x10^7$ PFU of the single and double recombinant sMVA-CoV2 vectors derived with FPV HP1.441 (sMVA-S/N hp and sMVA-N/S hp) or TROVAC (sMVA-S/N tv, sMVA-N/S tv, sMVA-S tv, sMVA-N tv) were evaluated for ADE effects. Neutralizing (1:5,000) and non-neutralizing (1:50,000) dilutions (as assayed on stably-transduced HEK293T cells expressing ACE2 (HEK/ACE2)) were evaluated to promote THP-1 monocyte infection by SARS-CoV-2 pseudovirus (pv) expressing luciferase. VSV-G pv was used as infection control. Relative light units (RLU) were measured in duplicates at 48 hours post infection Dotted lines represent the negative control (average relative light units [RLU] measured in cells in the absence of pv). Dashed lines represent the positive control (average RLU measured in cells in the absence of serum and in the presence of pv). 2-way ANOVA with Dunnett's multiple comparison test was used to compare each group and serum dilution to the mean RLU in the positive control. ns= not significant; *p<0.05.

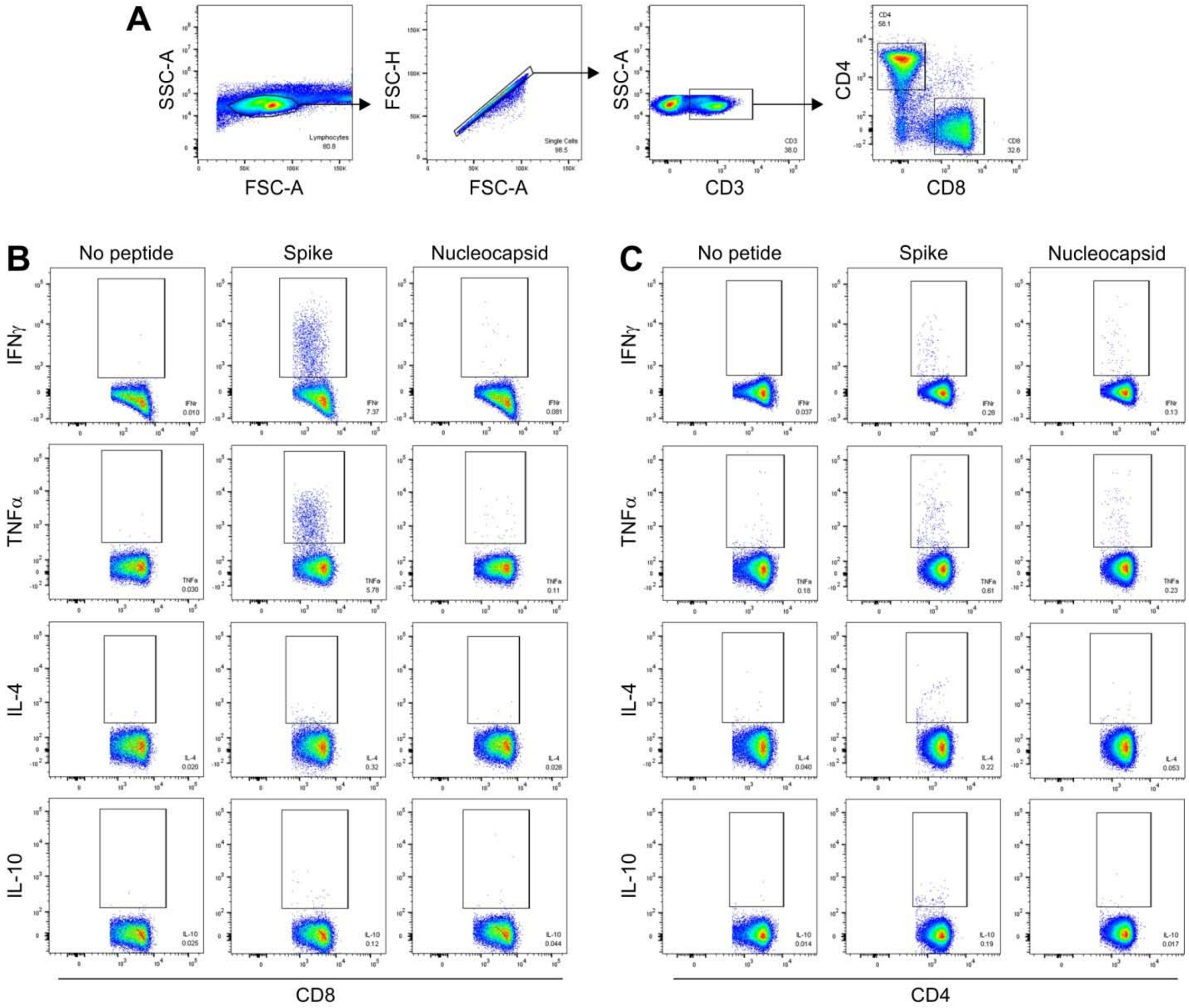


**Figure S7. Related to figure 6. Flow cytometry gating strategy. A)** Intracellular staining analysis of mouse splenocytes stimulated with S and N peptide libraries was performed using a hierarchical gating strategy that included lymphocytes>singlets>CD3$^{+}$ T-cells>CD4$^{+}$ T-cells and CD8$^{+}$ T-cells>Cytokine positive cells. **B-C)** Example of gating on cytokine-positive CD8$^{+}$ T-cells (B) and CD4$^{+}$ T-cells (C). Splenocytes of a mouse immunized with double recombinant vector sMVA-N/S were either left untreated (no peptide) or stimulated 16 hours with S or N peptide pools. Numbers in each dot plot indicate the percentage of cells in gated areas.

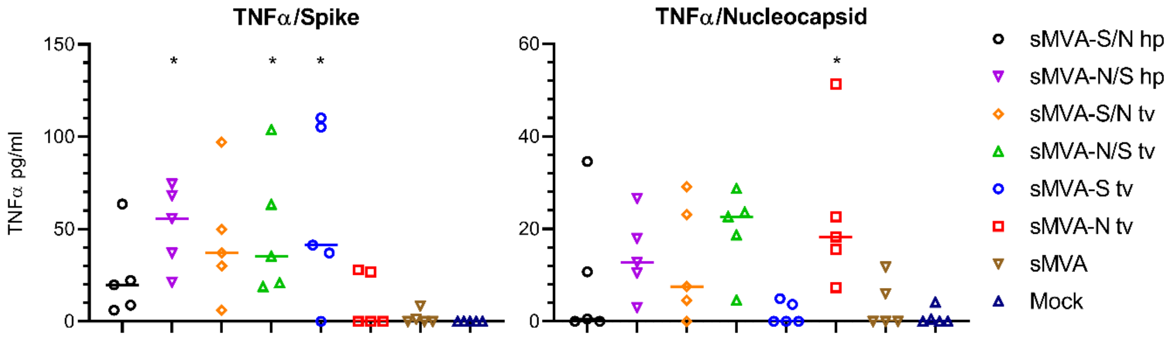


**Figure S8. Related to figure 6. TNFα secretion by T-cells of sMVA-CoV2-immunized mice.** Splenoctyes from Balb/c mice immunized with $5x10^7$ PFU of the single and double recombinant sMVA-CoV2 vectors derived with FPV HP1.441 (sMVA-S/N hp and sMVA-N/S hp) or TROVAC (sMVA-S/N tv, sMVA-N/S tv, sMVA-S tv, sMVA-N tv) were evaluated for TNFα secretion. Mouse splenocytes were stimulated with S or N peptide libraries and 48 hours later TNFα was measured by ELISA in cell culture supernatants. Amounts of TNFα quantified in unstimulated samples were subtracted from each peptide-stimulated sample. *p<0.05 compared to mock-immunized mice using one-way ANOVA with Dunnett's multiple comparison test.

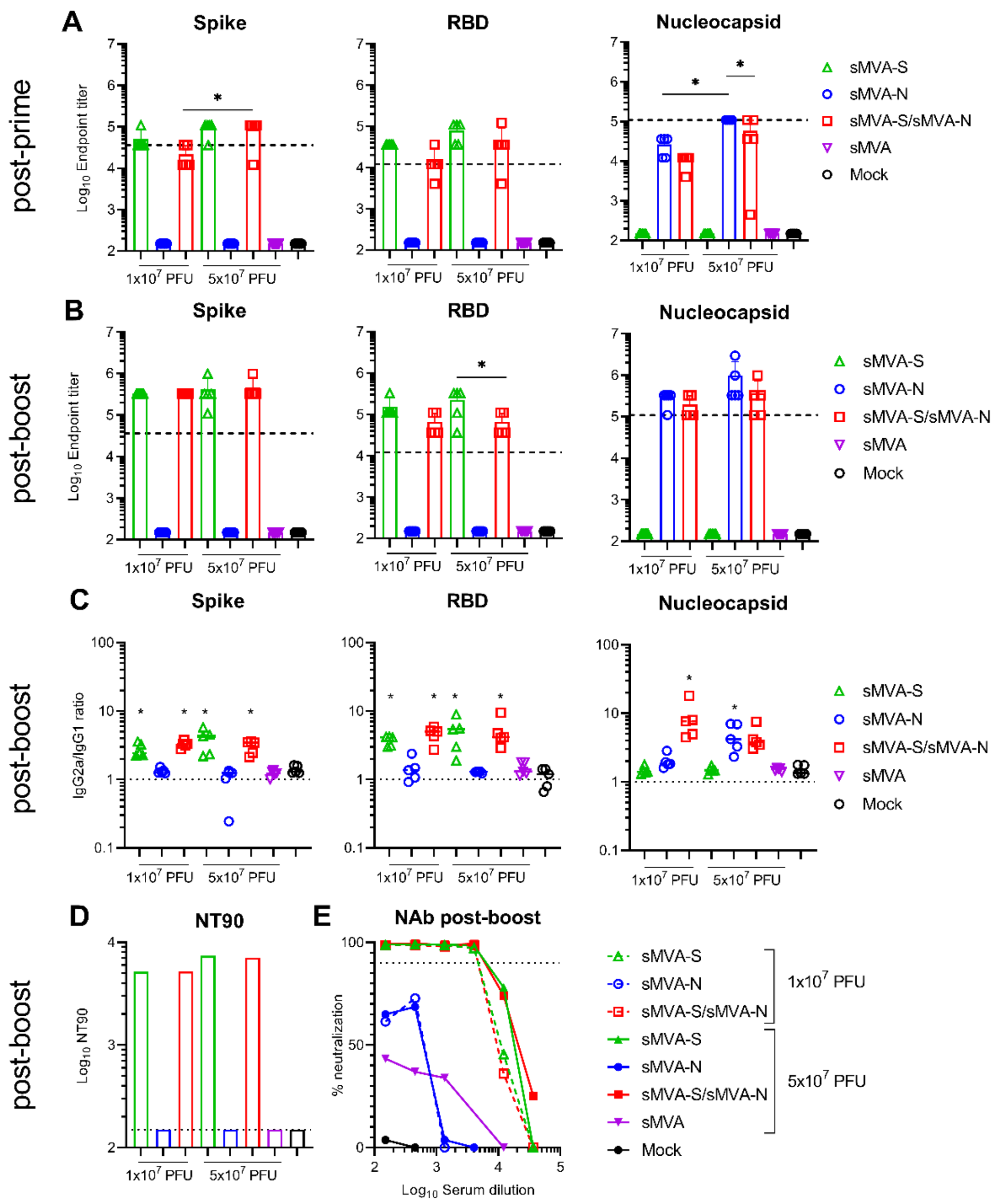


**Figure S9. Related to Figure 5. Humoral immune responses induced by sMVA-CoV2 vectors.** SARS-CoV-2-specific humoral immune responses were evaluated in mice immunized with the single recombinant vectors sMVA-S and sMVA-N alone or in combination. Balb/c mice (N=5) were immunized twice in three week interval with high ($5x10^7$ PFU) or low ($1x10^7$ PFU) dose of sMVA-S and sMVA-N. Co-immunization via the same immunization schedule with half of the high or low dose of each of the vaccine vectors was evaluated to assess SARS-CoV-2-specific immune stimulation to the S and N antigens by the vectors in combination. Mice immunized with empty sMVA vector or mock-immunized mice were used as controls. **A-B)** Binding antibodies. Antigen-specific binding antibodies to S, RBD, and N were determined after the first and second immunization by ELISA. Dashed lines indicate median binding antibody endpoint titers that were measured in convalescent human sera (Figure S4). One-way ANOVA with Tukey's multiple comparison test was used to compare differences between binding antibody end-point titers in mice immunized with different vaccine doses, and mice immunized with the vaccine vectors alone or combined. **C)** IgG2a/IgG1 isotype ratio. Ratio of IgG2a/IgG1 binding antibodies to S, RBD, and N was calculated after performing isotype-specific ELISA for the different antigens using post-boost serum from immunized mice. One-way ANOVA with Dunnett's multiple comparison test was used to compare each group mean to a ratio of 1 (balanced Th1/Th2 response). **D-E)** NAb titers. SARS-CoV-2-specific NAb responses were measured after the second immunization in pooled sera by neutralization assay using SARS-CoV-2 pseudovirus. Shown in D are the neutralizing antibody titers to prevent 90% infection of SARS-CoV-2 pseudovirus (NT90). Dotted baseline represents the minimum dilution included in the analysis. Groups with NT90<baseline are shown at baseline. E shows % neutralization measured using serial dilutions of pooled sera. Dotted line in E marks 90% neutralization. *$p<0.05$.

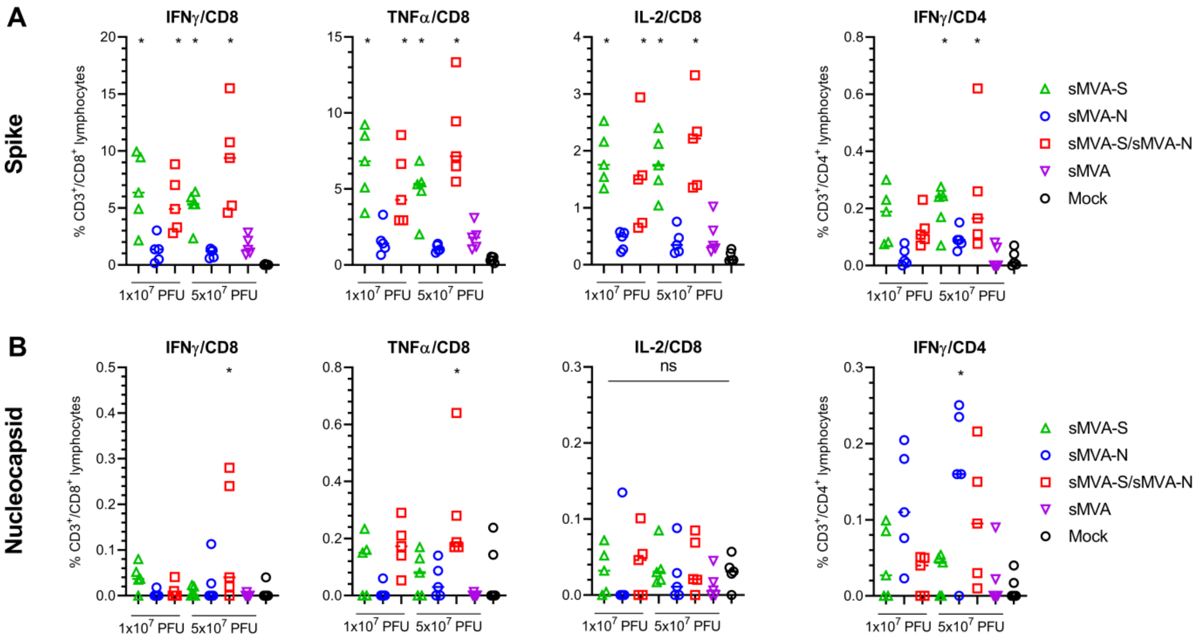


**Figure S10. Related to Figure 6. Cellular immune responses *In vivo* immunogenicity of sMVA-CoV2 vectors.** SARS-CoV-2-specific cellular immune responses were evaluated in mice immunized with the single recombinant vectors sMVA-S and sMVA-N alone or in combination. Balb/c mice (N=5) were immunized twice in three week interval with high ($5x10^7$ PFU) or low ($1x10^7$ PFU) dose of sMVA-S and sMVA-N. Co-immunization via the same immunization schedule with half of the high or low dose of each of the vaccine vectors was evaluated to assess SARS-CoV-2-specific immune stimulation to the S and N antigens by the vaccine vectors in combination. Mice immunized with empty sMVA vector or mock-immunized mice were used as controls. Antigen-specific CD8+ T cells expressing IFNγ, TNFα, and IL-2 and CD4+ T cell expressing IFNγ were evaluated by flow cytometry staining following *ex vivo* antigen stimulation using SARS-CoV-2-specific S and N peptide libraries. One-way ANOVA followed by Dunnett's multiple comparison test was used to compare each group mean to the mean in mock-immunized mice. *p<0.05.ns=not significant.

Table.S1

| Construct | Insert (insertion site) | Titer* |
| --- | --- | --- |
| sMVA hp | None | $6.8x10^{9}$ PFU/ml |
| sMVA tv1 | None | $4.1x10^{9}$ PFU/ml |
| sMVA tv2 | None | $2.3x10^{9}$ PFU/ml |
| wtMVA | None | $4.1x10^{9}$ PFU/ml |
| sMVA-S tv | Spike (Del3) | $4.3x10^{9}$ PFU/ml |
| sMVA-N tv | Nucleocapsid (Del3) | $1.0x10^{10}$ PFU/ml |
| sMVA-S/N hp | Spike (G1L), Nucleocapsid (Del3) | $8.8x10^{9}$ PFU/ml |
| sMVA-N/S hp | Nucleocapsid (Del2), Spike (Del3) | $2.3x10^{9}$ PFU/ml |
| sMVA-S/N tv | Spike (G1L), Nucleocapsid (Del3) | $8.8x10^{9}$ PFU/ml |
| sMVA-N/S tv | Nucleocapsid (Del2), Spike (Del3) | $8.4x10^{9}$ PFU/ml |

*Stocks were produced on CEF following infection (MOI 0.02) of 30x15cm dishes
